# Identification of the Polo-like kinase substrate required for homologous synapsis in *C. elegans*

**DOI:** 10.1101/2024.08.13.607834

**Authors:** Ariel L. Gold, Matthew E. Hurlock, Alicia M. Guevara, Lilah Y. Z. Isenberg, Yumi Kim

## Abstract

The synaptonemal complex (SC) is a zipper-like protein structure that aligns homologous chromosome pairs and regulates recombination during meiosis. Despite its conserved appearance and function, how synapsis occurs between chromosome axes remains elusive. Here, we demonstrate that Polo-like kinases (PLKs) phosphorylate a single conserved residue in the disordered C-terminal tails of two paralogous SC subunits, SYP-5 and SYP-6, to establish an electrostatic interface between the SC central region and chromosome axes in *C. elegans*. While SYP-5/6 phosphorylation is dispensable for the ability of SC proteins to self-assemble, local phosphorylation by PLKs at the pairing center is crucial for SC elongation between homologous chromosome axes. Additionally, SYP-5/6 phosphorylation is essential for asymmetric SC disassembly and proper PLK-2 localization after crossover designation, which drives chromosome remodeling required for homolog separation during meiosis I. This work identifies a key regulatory mechanism by which localized PLK activity mediates the SC-axis interaction through phosphorylation of SYP-5/6, coupling synapsis initiation to homolog pairing.

## INTRODUCTION

Accurate chromosome segregation during meiosis requires chromosomes to pair and undergo crossover recombination with their homologs in meiotic prophase. In most eukaryotes, homolog alignment is reinforced by synapsis, a process defined by the formation of a zipper-like protein assembly called the synaptonemal complex (SC). The SC is conventionally viewed as a tripartite structure consisting of two parallel chromosome axes that form between sister chromatids and a central region that connects the two (Page and Hawley, 2004). While the SC proteins can self-assemble to form aggregates known as “polycomplexes” outside chromosomes in diverse species (Moses, 1968; Goldstein, 1987; Zickler and Kleckner, 1999), the SC preferentially assembles between paired chromosome axes. The chromosome axis is composed of meiotic cohesins and, in most species, members of the HORMA domain protein family (Ur and Corbett, 2021). In budding yeast, plants, and mammals, HORMA domain proteins are depleted from axes upon SC assembly (Börner et al., 2008; Lambing et al., 2015; Wojtasz et al., 2009), and the changes in the axis composition enable the cell to coordinate synapsis with meiotic progression (Daniel et al., 2011; Kim et al., 2015). However, the interface between chromosome axes and the SC central region has not been clearly defined (Gordon and Rog, 2023).

The nematode *Caenorhabditis elegans* is an excellent model organism for investigating the molecular architecture and assembly mechanisms of the SC. Exhaustive genetic screens and biochemical purification have established a nearly complete list of proteins comprising the chromosome axis (Couteau et al., 2004; Couteau and Zetka, 2005; Martinez-Perez and Villeneuve, 2005; Goodyer et al., 2008; Wang et al., 2024; de Carvalho et al., 2008) and the SC central region in *C. elegans* (MacQueen et al., 2002; Colaiácovo et al., 2003; Smolikov et al., 2007, 2009; Hurlock et al., 2020; Zhang et al., 2020; Blundon et al., 2024). Soluble protein complexes representing the hierarchical assembly of the meiotic HORMA domain proteins (HIM-3, HTP-1, HTP-2, and HTP-3) and two other axis proteins (LAB-1 and LAB-2) have been reconstituted using purified proteins in vitro (Kim et al., 2014; Wang et al., 2024). A soluble SC building block comprising the six core subunits of the SC central region (SYP-1, SYP-2, SYP-3, SYP-4, SYP-5/6, and SKR-1/2) has also been reconstituted in vitro (Blundon et al., 2024), providing a foundation for the structural determination of the SC. Moreover, advances in super-resolution microscopy have provided valuable insights into the organization and structural changes within the SC during meiotic progression (Köhler et al., 2017, 2022; Woglar et al., 2020).

Before entering meiotic prophase, SYP/SKR proteins nucleate to form polycomplexes that exhibit liquid-like properties (Rog et al., 2017). SC materials then load at the pairing centers, specialized regions near one end of each chromosome crucial for homolog recognition, and rapidly extend along the chromosome length (Rog and Dernburg, 2015; MacQueen et al., 2005). These pairing centers function as signaling hubs, recruiting meiosis-specific Checkpoint kinase 2 (CHK-2) and Polo-like kinases (PLK-1 and PLK-2) to tether chromosomes to the cytoskeletal network and initiate meiotic chromosome dynamics (Harper et al., 2011; Labella et al., 2011; Kim et al., 2015; Sato et al., 2009). An additional PLK-docking site is generated on SYP-1 in early meiotic prophase (Sato-Carlton et al., 2018). However, PLKs are preferentially localized and constrained at the pairing center to ensure proper homolog pairing and synapsis (Brandt et al., 2020; Roelens et al., 2019; Zhang et al., 2012). Following crossover designation, PLK-2 relocates to the crossover site and the docking site on SYP-1 (Brandt et al., 2020; Sato-Carlton et al., 2018; Woglar and Villeneuve, 2018), stabilizing the SC in a chromosome-autonomous manner to prevent further DNA double-strand break formation (Machovina et al., 2016; Pattabiraman et al., 2017; Nadarajan et al., 2017).

In holocentric *C. elegans*, a single crossover event divides the chromosome into its short and long arms, ultimately defining the site of cohesin release during meiosis I (Altendorfer et al., 2020; Nabeshima et al., 2005; Ferrandiz et al., 2018; Rogers et al., 2002; Kaitna et al., 2002). One of the earliest events that exhibit asymmetry relative to the crossover site is the enrichment of PLK-2 on the short arm (Pattabiraman et al., 2017; Sato-Carlton et al., 2018; Brandt et al., 2020). Using the kinase activity, PLK-2 reinforces its own docking site on the SC short arm and drives axis remodeling and asymmetric SC disassembly (Harper et al., 2011; Sato-Carlton et al., 2018; Brandt et al., 2020). However, despite its central role in orchestrating SC assembly and disassembly in *C. elegans*, PLK substrates are largely unknown. While one residue in SYP-4 is known to be phosphorylated by PLKs, transitioning the SC to a more stable state upon crossover designation, the SC forms even without this phosphorylation (Nadarajan et al., 2017), leaving the PLK substrates required for synapsis elusive. We hypothesized that PLKs promote SC assembly by directly phosphorylating its core subunits. Therefore, we searched for residues conforming to the PLK consensus motifs (D/E-X-pS/T-Φ; Φ represents a hydrophobic residue) (Nakajima et al., 2003) within the SC central region proteins. In this work, we focus on two paralogous proteins, SYP-5 and SYP-6, which traverse the width of the SC in a head-to-head manner, with their N-termini positioned at the center and the C-termini facing the chromosome axes (Hurlock et al., 2020). We demonstrate that PLKs phosphorylate a single conserved residue at the disordered C-terminal tails of SYP-5/6 to establish the electrostatic interface between chromosome axes and the SC central region. This phosphorylation is essential not only for SC assembly but also for reinforcing the SC short-arm identity during disassembly, thereby ensuring faithful chromosome segregation during meiosis.

## RESULTS

### SYP-5 is phosphorylated at its C-terminal tail during meiotic prophase

SYP-5 and SYP-6 contain clusters of putative PLK phosphorylation sites in their disordered C-terminal tails. In particular, SYP-5 S541 and SYP-6 T685 are positioned near the end of the conserved C-terminal acidic patches (**Fig. 1 A**), which are crucial for SC assembly (Hurlock et al., 2020; Zhang et al., 2020). To determine whether these two residues are phosphorylated in vivo, we sought to raise polyclonal antibodies against the phosphopeptides surrounding SYP-5 S541 and SYP-6 T685. The affinity-purified SYP-5 pS541 antibody recognized the phosphorylated SYP-5 S541 peptide but not the non-phosphorylated version (**Fig. S1 A**), demonstrating its phosphospecificity. However, our efforts to obtain an antibody specific to SYP-6 pT685 were unsuccessful. Therefore, we focused on characterizing the phosphorylation status of SYP-5 S541, anticipating that SYP-6 would follow a similar pattern.

**Figure 1.**
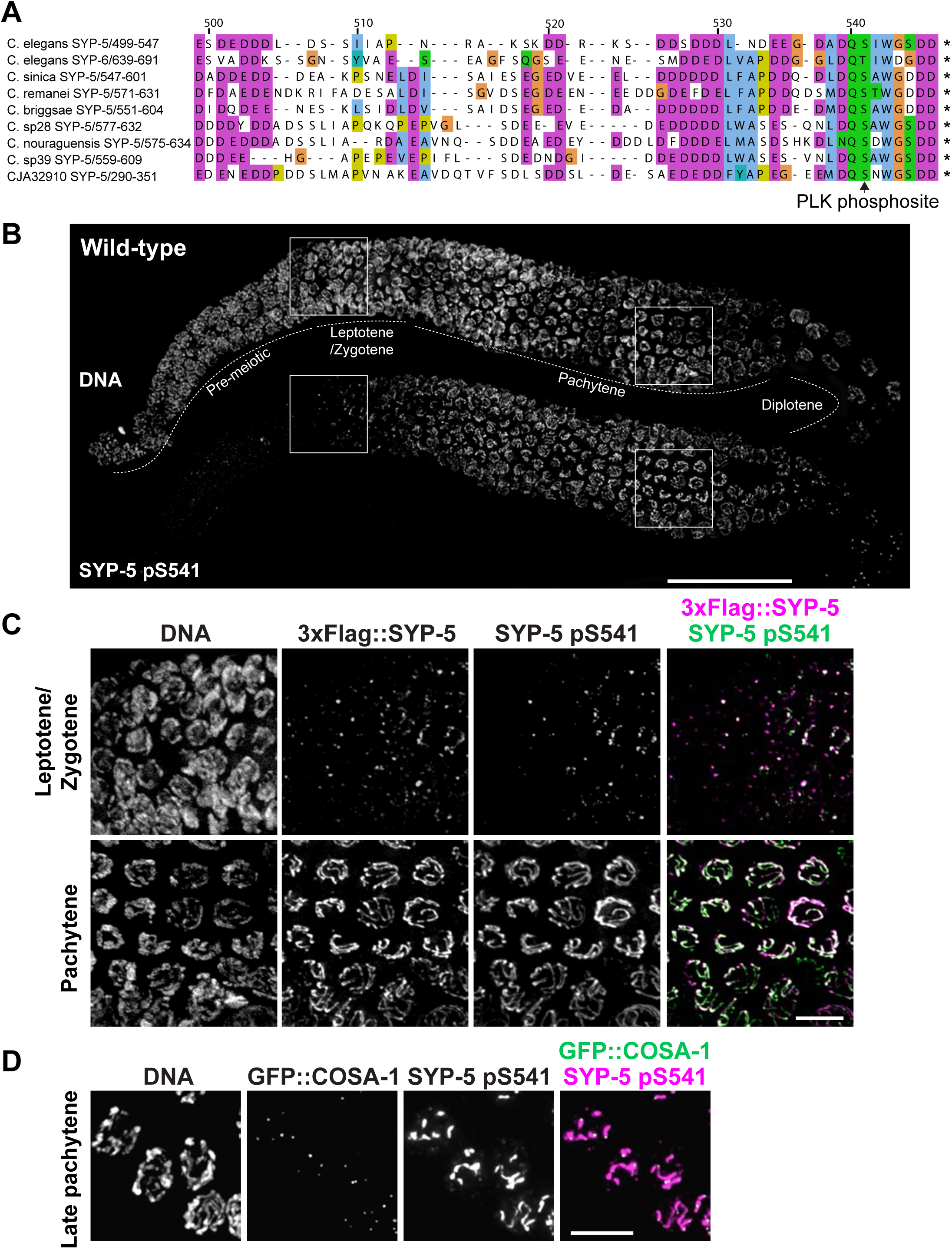
Phosphorylation of SYP-5 S541 during meiotic prophase. **(A)** Amino acid alignment of SYP-5/6 orthologs from *Caenorhabditis* species using the T-Coffee algorithm. The arrow indicates the PLK phosphorylation sites characterized in this study. **(B)** Composite projection images of a full-length gonad dissected from a worm strain expressing 3xFlag::SYP-5 and HA::SYP-6 are shown for DNA (top) and SYP-5 pS541 (bottom). Scale bar, 50 µm. **(C)** Zoomed-in images of leptotene/zygotene (upper panels) and pachytene nuclei (lower panels) as indicated by white boxes in (B). 3xFlag::SYP-5 (magenta) and SYP-5 pS541 (green) staining are shown. Scale bar, 10 µm. **(D)** Immunofluorescence images of late pachytene nuclei from animals expressing GFP::COSA-1. DNA, GFP::COSA-1 (green), and SYP-5 pS541 (magenta) staining are shown. Scale bar, 5 µm.

Immunofluorescence using the SYP-5 pS541 antibody revealed that SYP-5 is indeed phosphorylated at its C-terminal tail during meiotic prophase (**Fig. 1 B**). The phospho-signal for SYP-5 S541 was detected around meiotic entry at polycomplexes and along the SC in pachytene (**Fig. 1, B and C**). As meiosis progressed, SYP-5 S541 phosphorylation became restricted to one side of each crossover-designated site, marked by COSA-1 (Yokoo et al., 2012), in late pachytene (**Fig. 1 D**), representing the short arm of the SC. This localization aligns with the presence of PLK-2 on the SC in late pachytene (Sato-Carlton et al., 2018; Pattabiraman et al., 2017; Brandt et al., 2020; Nadarajan et al., 2017), supporting the hypothesis that PLKs might phosphorylate the C-terminal tails of SYP-5/6.

### Phosphorylation of SYP-5 at S541 requires both PLK-1 and PLK-2, but not the PLK-docking site on SYP-1

To demonstrate the phosphorylation of SYP-5 by PLKs, we performed in vitro kinase assays using recombinant 6His-MBP-SYP-5 and PLK-2 and mapped phosphorylation sites by mass spectrometry. Our analysis revealed robust phosphorylation of SYP-5 at multiple residues including S541 (**Fig. 2 A**). Additionally, the SYP-5 pS541 antibody recognized recombinant 6His-MBP-SYP-5 after incubation with PLK-2 (**Fig. 2 B**), confirming the phosphorylation of SYP-5 S541 by PLK-2 in vitro. To determine whether PLKs are responsible for SYP-5 phosphorylation in vivo, we examined the phosphorylation status at SYP-5 S541 in wild-type, *plk-2,* and *plk-2; plk-1(RNAi)* animals by immunofluorescence. Due to partial redundancy between PLK-2 and its mitotic paralog PLK-1 (Harper et al., 2011; Labella et al., 2011), we knocked down *plk-1* via RNAi in *plk-2* mutants to deplete both PLKs. While SYP-5 S541 phosphorylation was robustly detected along the SC in wild-type and *plk-2* animals, it was abolished in *plk-2; plk-1(RNAi)* mutants (**Fig. 2 C**). Thus, SYP-5 S541 phosphorylation depends on both PLK-1 and PLK-2 in vivo. We also determined whether the recruitment of PLKs to the SC is necessary for SYP-5 phosphorylation by staining for SYP-5 pS541 in *syp-1^T452A^* mutants, which are defective in PLK docking (Sato-Carlton et al., 2018; Brandt et al., 2020). Interestingly, SYP-5 phosphorylation was still detected on the SC in *syp-1^T452A^* animals **(Fig. 2 C)**, indicating that SYP-5 phosphorylation does not require the PLK-docking site on the SC.

**Figure 2.**
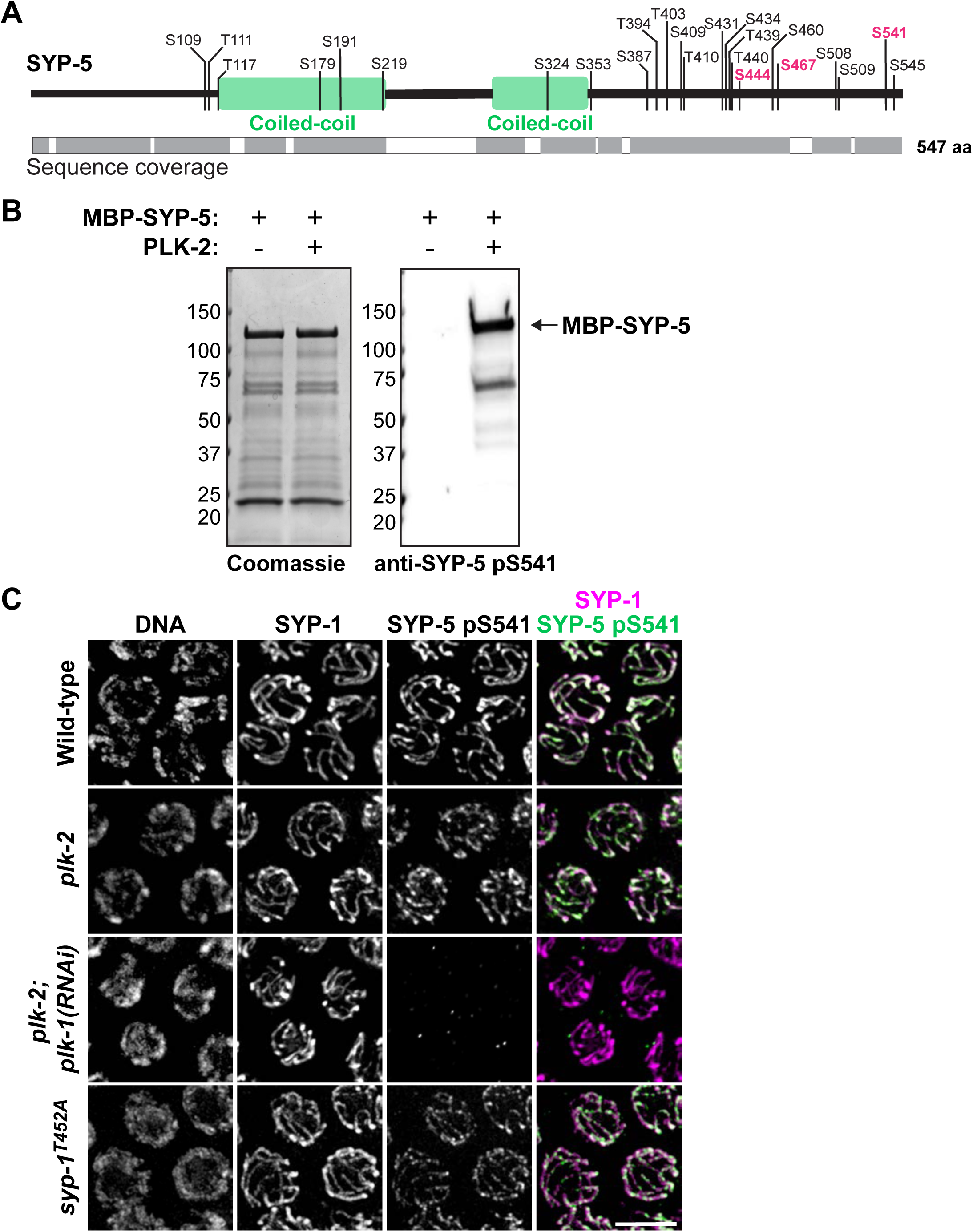
Phosphorylation of SYP-5 S541 requires both PLK-1 and PLK-2, but not the PLK-docking site on SYP-1. **(A)** Schematic showing in vitro PLK-2 phosphorylation sites on SYP-5, with sequence coverage from mass spectrometry analysis indicated by gray blocks. Conserved residues that conform to the PLK consensus motif are highlighted in magenta. **(B)** In vitro kinase assays using recombinant 6His-MBP-SYP-5 incubated with or without PLK-2. Coomassie staining of the purified proteins (left) and western blot for SYP-5 pS541 (right) are shown. 6His-MBP-SYP-5 is indicated by arrow. **(C)** Immunofluorescence images of pachytene nuclei from wild-type, *plk-2(tm1395), plk-2(tm1395); plk-1(RNAi*), and *syp-1^T452A^* animals, stained for SYP-1 (magenta), and SYP-5 pS541 (green). Scale bar, 5 µm.

### Phosphorylation of SYP-5 S541 and SYP-6 T685 is essential for SC assembly

Next, we investigated the significance of SYP-5/6 C-terminal phosphorylation by introducing phosphorylation-defective mutations at SYP-5 S541 and SYP-6 T685 using CRISPR. The self-progeny of hermaphrodites homozygous for the *syp-5^S541A^ syp-6^T685A^*mutations exhibited a dramatic reduction in egg viability (49% vs. 91% in the wild-type control) (**Fig. S2 A**), and 9% of surviving progeny were males, in contrast to 2% observed in control (**Fig. S2 B**). In *C. elegans*, males arise from X-chromosome nondisjunction during meiosis (Hodgkin et al., 1979), implying errors in meiotic chromosome segregation in *syp-5^S541A^ syp-6^T685A^* mutants. We note a higher embryonic lethality and percentage of males in our control strain than N2 (100% egg viability; 0.2% males), which is likely due to the epitopes tagged to SYP-5/6 (*3xflag::syp-5 ha::syp-6*) (Hurlock et al., 2020).

*syp-5^S541A^ syp-6^T685A^*animals were proficient in homolog pairing, as assessed by the X chromosome pairing center protein, HIM-8 (Phillips et al., 2005) (**Fig. S2 C**). However, these animals exhibited severe defects in SC assembly, with most cells displaying either polycomplexes or only partial SC stretches in the germline (**Fig. 3 A**). Importantly, the phospho-signal at SYP-5 S541 was eliminated in the *syp-5^S541A^ syp-6^T685A^*mutants (**Fig. S1 B**). Failures in synapsis cause delays in meiotic progression by extending the kinase activity of CHK-2 (Kim et al., 2015; Stamper et al., 2013; Rosu et al., 2013). Consistent with this, we found that the “CHK-2 active zone” is substantially prolonged in *syp-5^S541A^ syp-6^T685A^* mutants (73% vs. 43% in the wild-type control), as examined by CHK-2-mediated phosphorylation of the pairing center proteins (HIM-8 and ZIMs) (Kim et al., 2015) (**Fig. S2, D and E**). Additionally, while control animals displayed six crossover-designated COSA-1 foci in late pachytene (Yokoo et al., 2012), the average number of COSA-1 foci in *syp-5^S541A^ syp-6^T685A^* mutants was reduced to 4.2, ranging from one to seven (**Fig. S2, F and G**), reflecting the role of the SC in both promoting and limiting crossovers (Hayashi et al., 2010; Libuda et al., 2013). Together, these results demonstrate that PLK-mediated phosphorylation at the single conserved residue in the C-termini of SYP-5/6 is essential for robust SC assembly.

**Figure 3.**
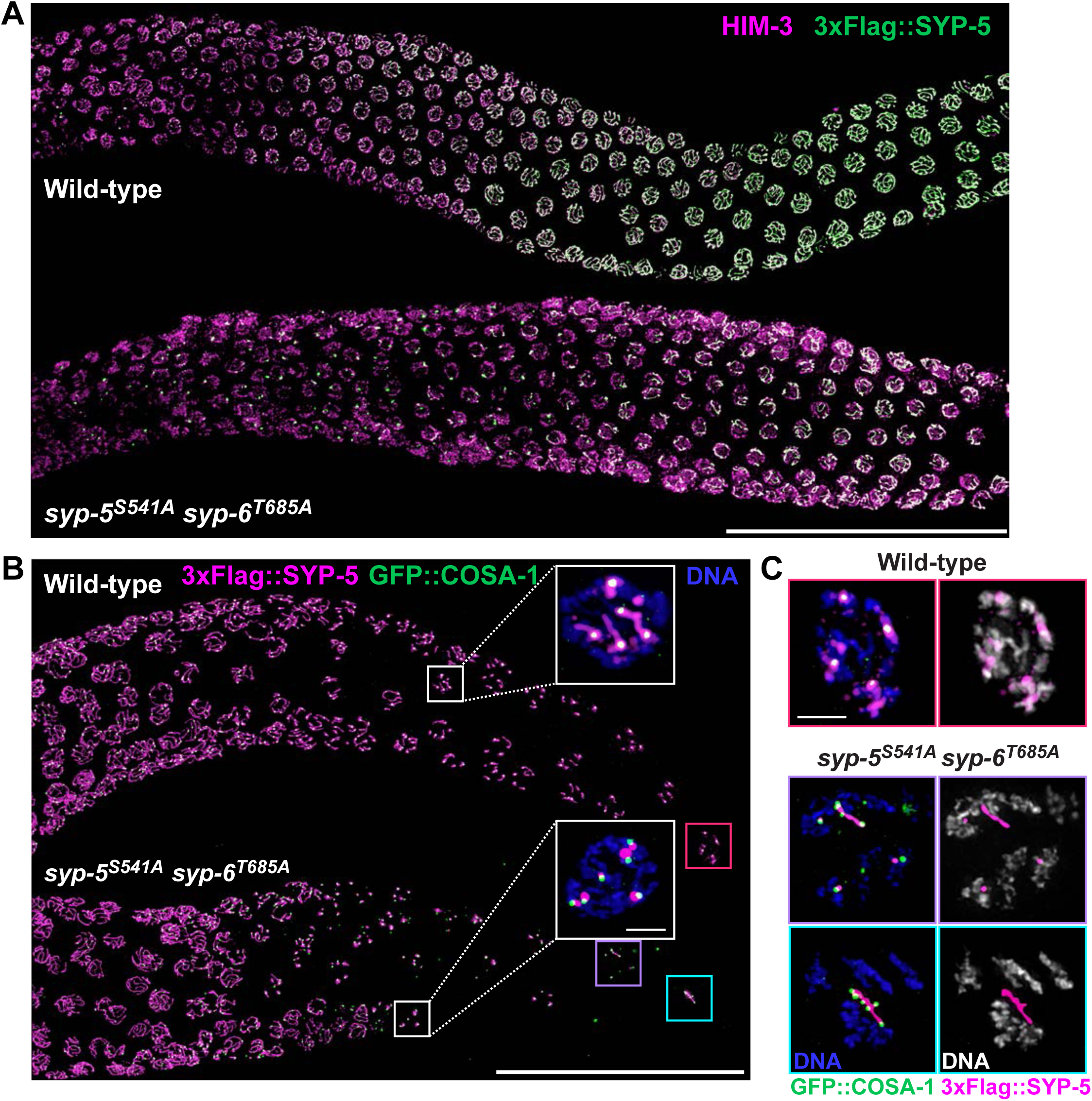
Phosphorylation of SYP-5 S541 and SYP-6 T685 is critical for proper SC assembly and disassembly. **(A)** Composite immunofluorescence images of full-length gonads from wild-type and *syp-5^S541A^ syp-6^T685A^* mutant animals stained for HIM-3 (magenta) and 3xFlag::SYP-5 (green). Scale bar, 50 µm. **(B)** Composite immunofluorescence images of late pachytene and diplotene region from wild-type and *syp-5^S541A^ syp-6^T685A^* animals showing 3xFlag::SYP-5 (magenta) and GFP::COSA-1 (green) staining. Scale bar, 50 µm. Insets show zoomed-in views of the boxed regions with DNA staining (blue). Scale bar, 1 µm. **(C)** Zoomed-in views of the boxed regions in diplotene as indicated by colored boxes in (B). DNA (blue), SYP-5 (magenta), and GFP::COSA-1 (green) staining are shown. Scale bar, 1 µm.

### Phosphomimetic mutations at the C-termini of SYP-5 and SYP-6 also lead to synapsis defects

We also mutated SYP-5 S541 and SYP-6 T685 to aspartic acids (D) to determine whether introducing negative charges at the SYP-5/6 C-termini is necessary for SC assembly. The self-progeny of hermaphrodites carrying the *syp-5^S541D^ syp-6^T685D^* mutation exhibited a marked decrease in egg viability (48% vs. 91% in control) and an increase in the percentage of male progeny (8% vs. 2% in control) (**Fig. S3, A and B**), indicating errors in meiotic chromosome segregation. Interestingly, *syp-5^S541D^ syp-6^T685D^*mutants also displayed severe defects in SC assembly (**Fig. S3 C**), similar to the phenotypes observed in *syp-5^S541A^ syp-6^T685A^* mutants. As a result, the CHK-2 active zone was longer in *syp-5^S541D^ syp-6^T685D^* mutants (74%), compared to 43% in control (**Fig. S3, D and E**). These data suggest that introducing negative charges to the C-terminal tails of SYP-5/6 is insufficient to promote synapsis. Alternatively, amino acid substitutions may not accurately recapitulate the effects of phosphorylation, and attaching phosphate groups to these residues is critical for SC assembly.

### Phosphorylation of SYP-5 S541 and SYP-6 T685 is essential for retaining SC proteins on chromosome axes after crossover designation

Further analyses of *syp-5^S541A^ syp-6^T685A^*and *syp-5^S541D^ syp-6^T685D^* mutants revealed that phosphorylation of the C-terminal tails of SYP-5/6 is necessary for proper SC disassembly. In the control strain, the SC central region disassembled from the long arm relative to the crossover-designated site and was retained on the short arm (**Fig. 3 B**), as previously described (Nabeshima et al., 2005; Martinez-Perez et al., 2008). However, in both *syp-5^S541A^ syp-6^T685A^* and *syp-5^S541D^ syp-6^T685D^* mutants, the SC proteins were prematurely lost from chromosome arms and appeared as puncta at crossover-designated sites (**Fig. 3 B and S4 A**). Strikingly, SC proteins frequently dissociated from condensing chromosomes in diplotene and formed elongated polycomplexes that associated with multiple COSA-1 foci in *syp-5^S541A^ syp-6^T685A^* (42% of the gonads, n=12) and *syp-5^S541D^ syp-6^T685D^* mutants (38% of the gonads, n=12) (**Fig. 3 C and S4 A**).

We hypothesized that phosphorylation of the SYP-5/6 C-termini is essential for maintaining the association of SC proteins with chromosome arms after crossover designation and that SC proteins primarily associate with crossover-designated sites in the absence of SYP-5/6 phosphorylation. To test this hypothesis, we combined the *syp-5^S541A^ syp-6^T685A^* mutations with a null allele of *cosa-1* essential for crossover designation (Yokoo et al., 2012). In *cosa-1* mutants, the SC and phospho-signal for SYP-5 S541 persisted on chromosome axes until diplotene and diakinesis (**Fig. S4, B and C**). By contrast, in *syp-5^S541A^ syp-6^T685A^; cosa-1* animals, the SC proteins showed abrupt dissociation from condensing chromosomes and were instead found in nucleoplasmic aggregates (**Fig. S4 B**). Thus, phosphorylation of the SYP-5/6 C-termini is essential for retaining the SC central region proteins on the short arm during SC disassembly.

### Phosphorylation of SYP-5/6 C-termini is required for asymmetric partitioning of bivalents after crossover designation

After crossover designation, PLK-2 becomes enriched on the SC short arm via its docking site on SYP-1 (Sato-Carlton et al., 2018; Brandt et al., 2020). However, in *syp-5^S541A^ syp-6^T685A^* mutants, PLK-2::mRuby dissociates from chromosomes, accumulating instead on nucleoplasmic SC aggregates (**Fig. 4 A**). The displacement of PLK-2 led to an aberrant pattern of chromosome remodeling necessary for two-step cohesin loss during meiotic divisions. Notably, two paralogous HORMA domain proteins, HTP-1 and HTP-2, which are normally restricted to the long arm (Martinez-Perez et al., 2008), were observed on all chromosome arms in *syp-5^S541A^ syp-6^T685A^* mutants (**Fig. 4 B**). In the wild-type control, HTP-1 and HTP-2, along with the axis protein LAB-1, recruit protein phosphatase 1 (PP1) to counteract phosphorylation at histone H3 T3 on the long arm (Ferrandiz et al., 2018; de Carvalho et al., 2008), which serves as a docking site for the chromosome passenger complex necessary for releasing sister cohesion during meiosis I (Kaitna et al., 2002; Rogers et al., 2002; Kelly et al., 2010). In *syp-5^S541A^ syp-6^T685A^* mutants, however, the phospho-mark for histone H3 T3 was lost from chromosomes (**Fig. 4 C**), consistent with the persistent presence of HTP-1/2 and phosphatase activity along all chromosome arms. Thus, phosphorylation of the SYP-5/6 C-terminal tails is essential for establishing the SC short-arm identity and reinforcing the recruitment of PLK-2 to meiotic chromosomes.

**Figure 4.**
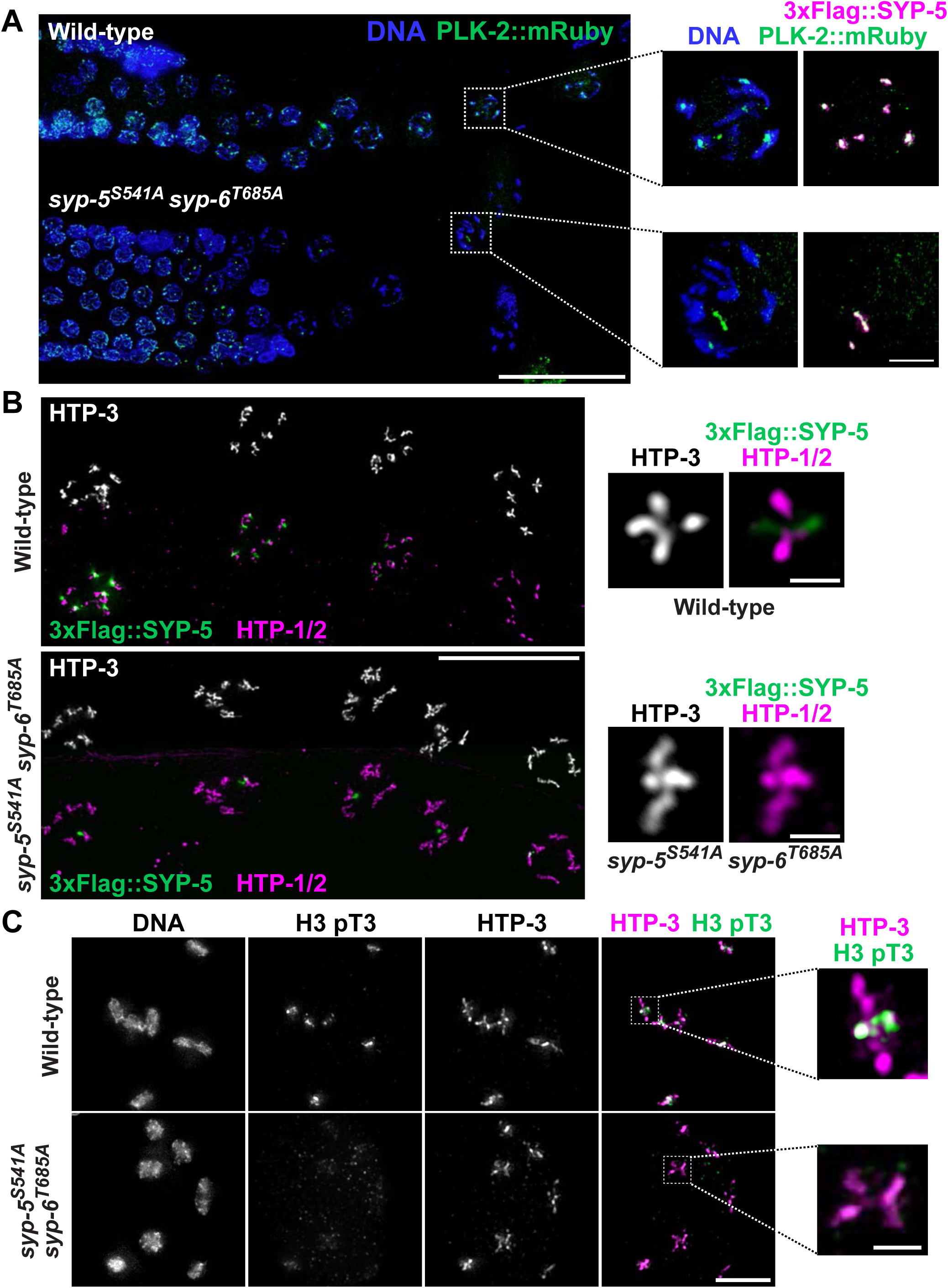
Phosphorylation of SYP-5/6 is required for proper PLK-2 localization and chromosome remodeling in late meiotic prophase. **(A)** Composite immunofluorescence images of wild-type and *syp-5^S541A^ syp-6^T685A^* animals showing DNA (blue) and PLK-2::mRuby (green) staining. Scale bar, 25 µm. Insets show zoomed-in views of the boxed regions with 3xFlag::SYP-5 (magenta) staining. Scale bar, 1 µm. **(B)** Immunofluorescence images of diakinesis nuclei from wild-type and *syp-5^S541A^ syp-6^T685A^* animals showing HTP-3, 3xFlag::SYP-5 (green), and HTP-1/2 (magenta). Scale bar, 20 µm. Insets show representative bivalents from indicated genotypes showing HTP-3, HTP-1/2 (magenta), and SYP-5 (green) staining. Scale bar, 1 µm. **(C)** Immunofluorescence images of diakinesis nuclei from wild-type and *syp-5^S541A^ syp-6^T685A^* animals showing DNA, histone H3 pT3 (green), and HTP-3 (magenta) staining. Scale bar, 5 µm. Insets show zoomed-in views of the boxed regions. Scale bar, 1 µm.

### Conserved tryptophan residues in SYP-5/6 C-termini are essential for proper SC assembly and disassembly, independent of SYP-5/6 phosphorylation by PLKs

Intriguingly, the PLK phosphorylation site within the SYP-5/6 C-termini is adjacent to a highly conserved tryptophan residue (SYP-5 W543 and SYP-6 W687), surrounded by negatively charged regions (**Fig. 1 A and Fig. 5, A and B**). The indole side chain of tryptophan often serves as a π donor to cations, providing a strong attractive force crucial for ligand binding and protein-protein interactions (Gallivan and Dougherty, 1999; Dougherty, 1996). We reasoned that this putative cation-π interaction could contribute to synapsis in conjunction with the charge-charge interactions provided by the phosphorylation of the acidic C-termini of SYP-5/6. To determine the significance of these tryptophan residues, we mutated SYP-5 W543 and SYP-6 W687 to alanines using CRISPR. Indeed, *syp-5^W543A^ syp-6^W687A^* animals exhibited severe defects in SC assembly and disassembly (**Fig. 5 C and S5 A**), similar to the phenotypes observed in *syp-5/6* mutants defective in phosphorylation (**Fig. 3**), though with less frequent nucleoplasmic SC aggregates in diplotene (17% of gonads, n=6). Notably, the phospho-signal for SYP-5 S541 was still detected in *syp-5^W543A^ syp-6^W687A^* mutants (**Fig. S5 B**), indicating that the tryptophan residues are crucial for SC assembly, independent of the phosphorylation at the SYP-5/6 C-termini.

**Figure 5.**
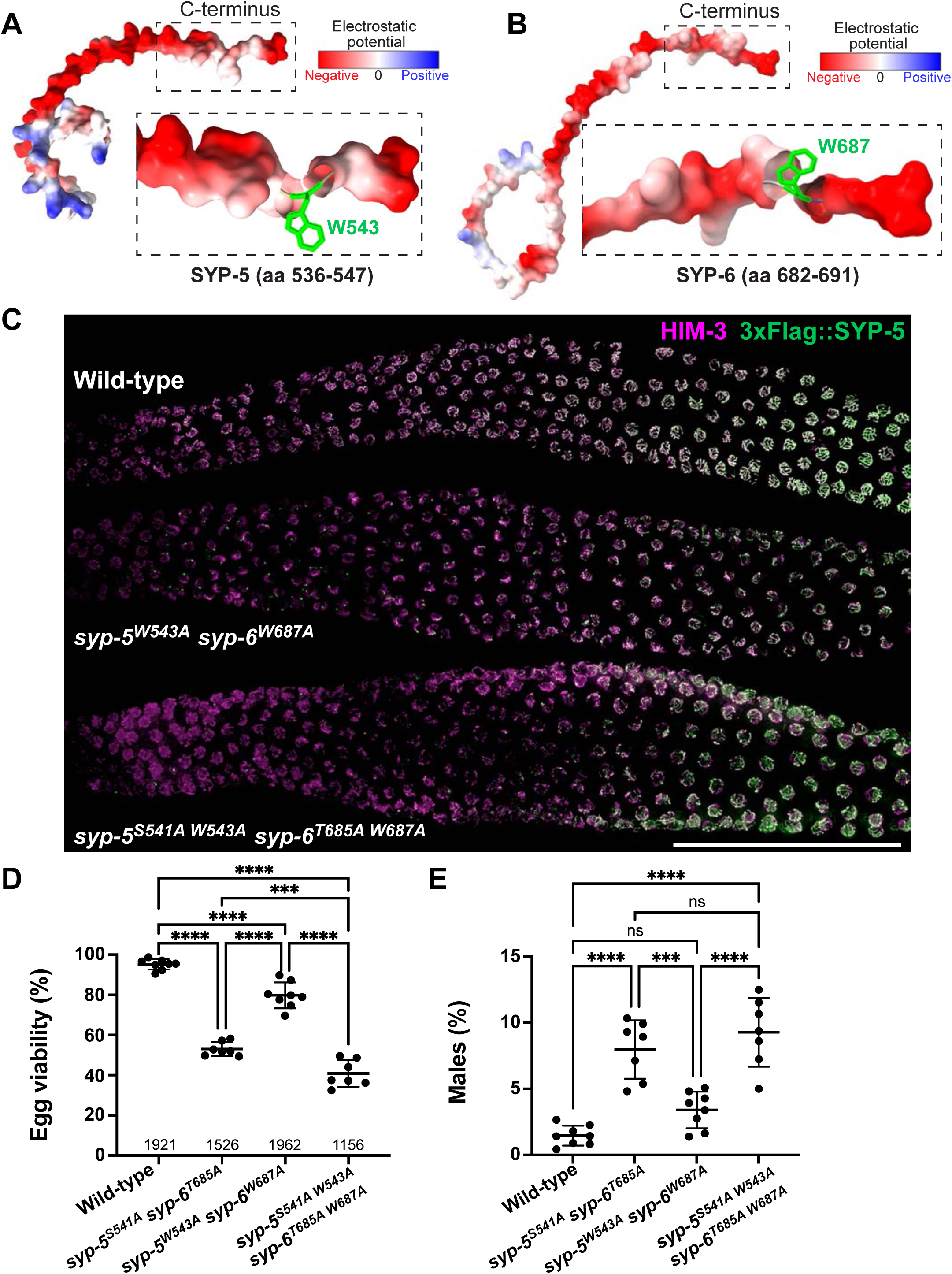
Conserved tryptophan residues in SYP-5/6 C-termini are essential for proper SC assembly. **(A-B)** Electrostatic potential of the C-terminal tails of SYP-5 (aa 536-547) (A) and SYP-6 (aa 682-691) (B) using AlphaFold models (AF-Q9N3E8-F1 and AF-A0A7R9SUK8-F1). Conserved tryptophan residues (SYP-5 W543 and SYP-6 W687) are labeled in green. **(C)** Composite immunofluorescence images of dissected gonads from wild-type, *syp-5^W543A^ syp-6^W687A^*, and *syp-5^S541A^ ^W543A^ syp-6^T685A^ ^W687A^* animals showing HIM-3 (magenta) and 3xFlag::SYP-5 (green). Scale bar, 50 µm. **(D and E)** Graphs showing the percent viable and male self-progeny from wild-type (n=1921), *syp-5^S541A^ syp-6^T685A^* (n=1526), *syp-5^W543A^ syp-6^W687A^* (n=1962), and *syp-5^S541A, W543A^ syp-6^T685A, W687A^* hermaphrodites (n=1156). ****, p<0.0001 by unpaired t-test.

We next determined whether mutating the tryptophan residues exacerbates the SC defects observed in *syp-5/6* phospho-mutants. Hermaphrodites carrying the *syp-5^S541A^ ^W543A^ syp-6^T685A^ ^W687A^* mutations produced an average of 41% viable self-progeny, with 9% of them being males, which are lower than the percentages observed in *syp-5^S541A^ syp-6^T685A^* (53%) or *syp-5^W543A^ syp-6^W687A^* mutants (80%) (**Fig. 5, D and E**). These data suggest that PLK-mediated phosphorylation and the putative cation-π interaction independently contribute to meiotic fidelity. Immunofluorescence analysis revealed similarly severe SC defects in *syp-5^S541A^ ^W543A^ syp-6^T685A^ ^W687A^* mutants (**Fig. 5 C**), but a higher frequency of nucleoplasmic SC aggregates in diplotene (75% of the gonads, n=8) (**Fig. S5 A**), potentially explaining the additive effect of the tryptophan mutations on egg viability in *syp-5/6* phospho-mutants. Additionally, western blot analysis showed that SYP-5 levels were unchanged by any of the mutations we generated in the C-terminal tails of SYP-5/6 (**Fig. S5 C**), indicating that the SC defects are not due to altered SYP-5/6 expression.

### Phosphorylation and tryptophan residues in SYP-5/6 C-termini provide affinity between the SC central region and meiotic HORMA domain proteins

We hypothesized that PLK-mediated phosphorylation and the aromatic side chain of tryptophan residues at the SYP-5/6 C-termini provide affinity between the SC central region and chromosome axes, coordinating the assembly and disassembly of the SC during meiotic progression. The interaction between the SC central region and axis components is transient and not readily detected by conventional immunoprecipitation (Hurlock et al., 2020; Zhang et al., 2020; Blundon et al., 2024). However, HORMA domain proteins are recruited to SC polycomplexes when meiosis-specific cohesin complexes are absent in *C. elegans* (Pasierbek et al., 2001; Severson et al., 2009). Therefore, we used a triple cohesin mutant that lacks all meiotic kleisins (*rec-8; coh-3 coh-4*) and examined the recruitment of HTP-3 and HIM-3 to polycomplexes. As expected, HIM-3 and HTP-3 were found at polycomplexes in our control strain (**Fig. 6 A**). By contrast, when the null alleles of *rec-8, coh-3, and coh-4* were combined with either *syp-5^S541A^ syp-6^T685A^*, *syp-5^S541D^ syp-6^T685D^*, or *syp-5^W543A^ syp-6^W687A^* mutations, HIM-3 and HTP-3 failed to localize to spherical polycomplexes (**Fig. 6 A**). Thus, while SYP-5/6 phosphorylation and the tryptophan residues are dispensable for the overall ability of SC proteins to form polycomplexes, they are essential for establishing the interface between the SC central region and meiotic HORMA domain proteins.

**Figure 6.**
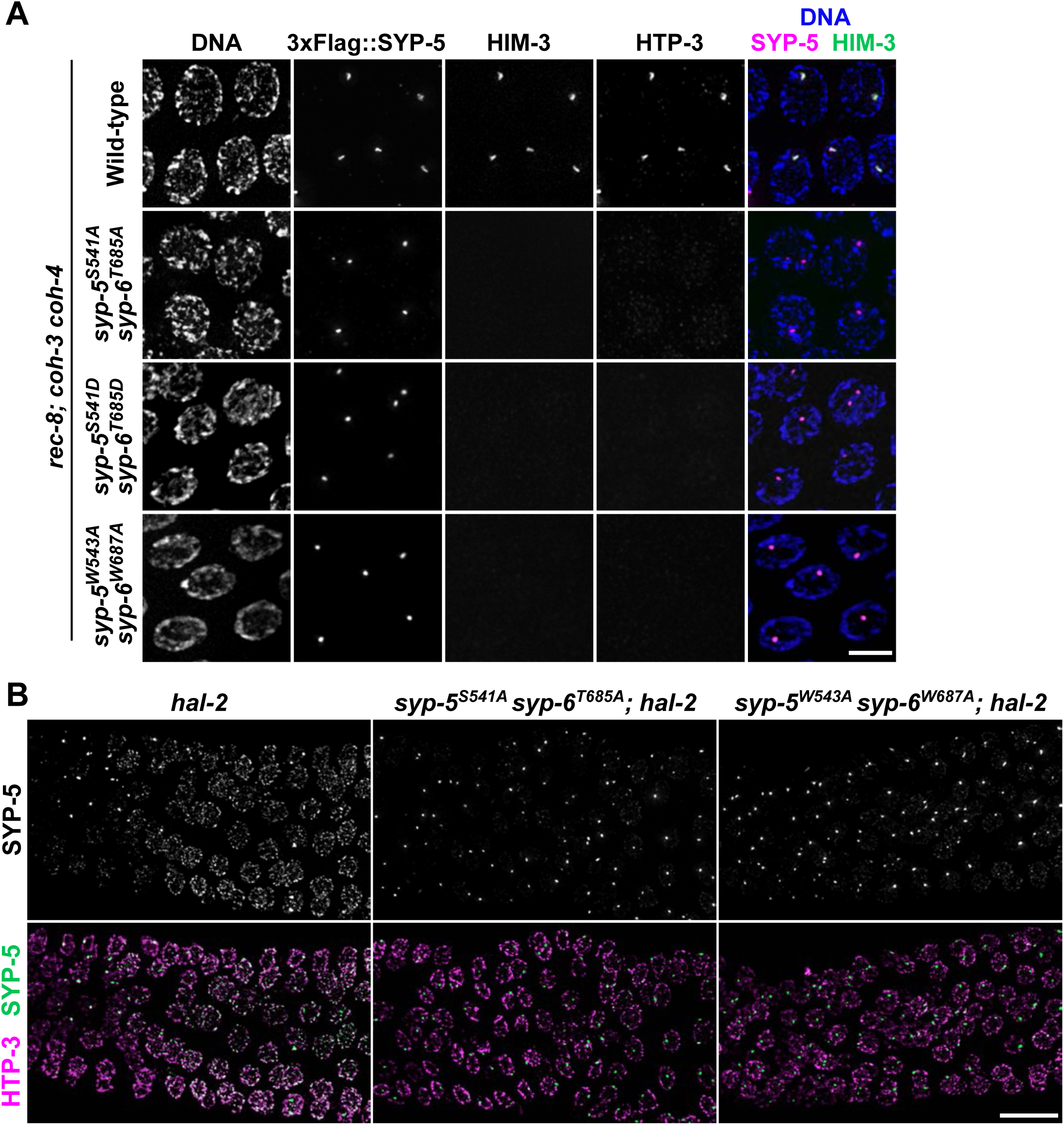
Phosphorylation of SYP-5/6 C-termini mediate interactions between the SC central region and meiotic HORMA domain proteins. **(A)** Immunofluorescence images of pachytene nuclei from indicated genotypes lacking meiotic kleisin subunits (*rec-8; coh-3 coh-4*). DNA (blue), 3xFlag::SYP-5 (magenta), HIM-3 (green), and HTP-3 staining are shown. Scale bar, 5 µm. **(B)** Composite immunofluorescence images of early pachytene nuclei dissected from indicated genotypes showing SYP-5 (green) and HTP-3 (magenta) staining. Scale bar, 10 µm.

Finally, we investigated whether the C-termini of SYP-5/6 are the major PLK substrates responsible for the affinity between the SC central region and chromosome axes. During early meiotic prophase, PLK activity is restricted to the pairing centers by the HAL-2/3 complex. However, in *hal-2/3* mutants, unleashed PLK activity in the nucleoplasm leads to premature association of SC proteins with unpaired chromosome axes (Zhang et al., 2012; Roelens et al., 2019). Interestingly, the loading of SYP-5 to chromosome axes in *hal-2* mutants was prevented by *syp-5^S541A^ syp-6^T685A^* or *syp-5^W543A^ syp-6^W687A^*mutations, resulting in SYP-5 detected as polycomplexes throughout the germline (**Fig. 6 B**). These data suggest that the C-terminal tails of SYP-5/6 are the primary PLK substrates crucial for SC loading and also highlight the necessity of tightly controlled phosphorylation of SYP-5/6.

## DISCUSSION

Here, we identify a single conserved residue in the disordered C-terminal tails of SYP-5/6 as the primary PLK substrate essential for homologous synapsis in *C. elegans* meiosis. While phosphorylation of SYP-5/6 is dispensable for the self-assembly of SC subunits, it is crucial for establishing the interface between the SC central region and chromosome axes. Furthermore, SYP-5/6 phosphorylation reinforces the identity of the SC short arm and PLK-2 localization following crossover designation, thereby ensuring chromosome remodeling required for proper homolog separation during meiosis I (**Fig. 7**).

**Figure 7.**
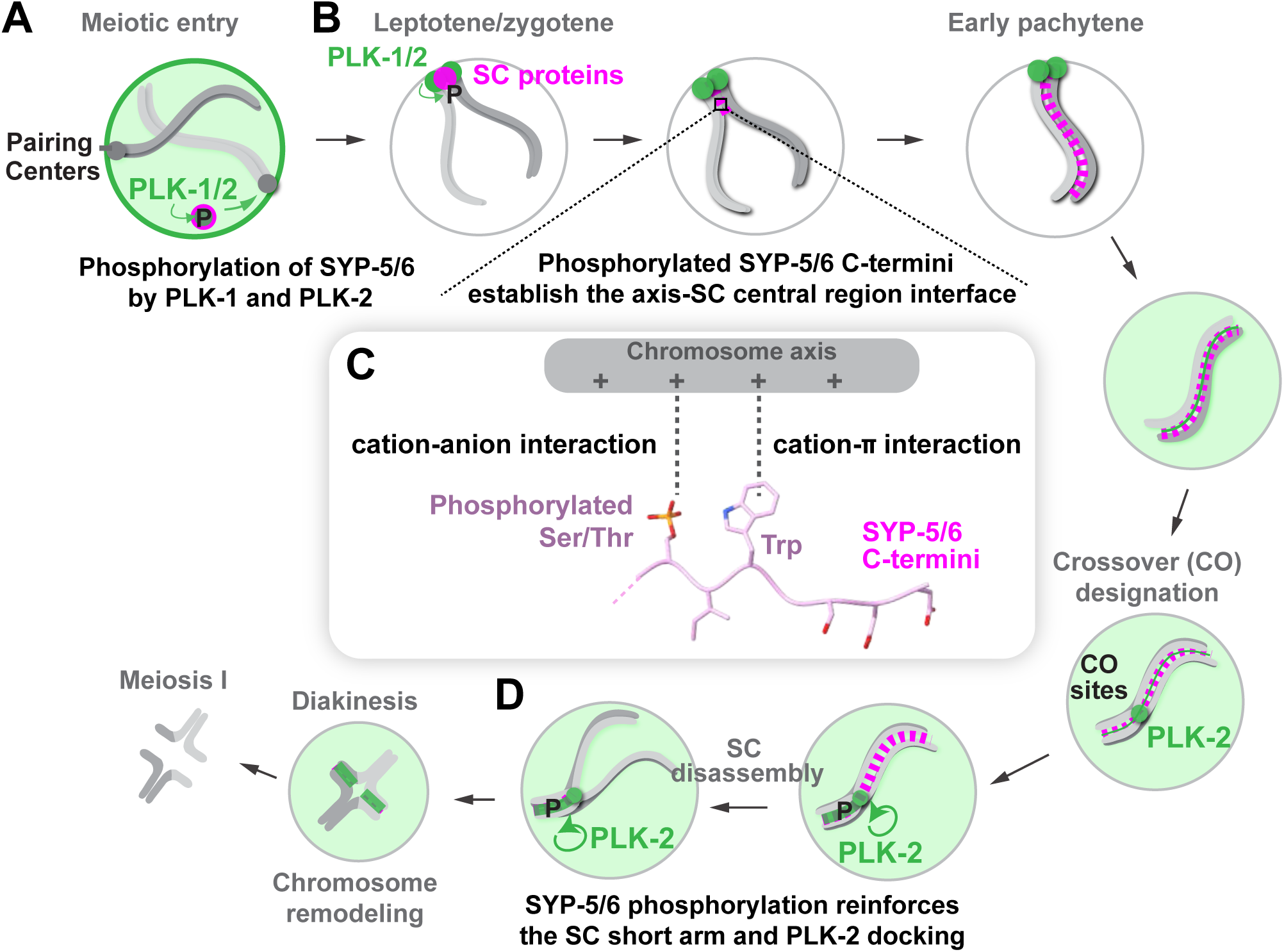
Model for PLK-mediated regulation of SC assembly and disassembly via phosphorylation of SYP-5/6 C-termini in *C. elegans*. **(A)** PLK-dependent phosphorylation of the C-termini of SYP-5/6 occurs just prior to meiotic entry. **(B)** Localized phosphorylation of SYP-5/6 by PLKs at the pairing centers links homolog pairing with the initiation of synapsis. **(C)** Phosphorylation of the SYP-5/6 C-termini establishes an electrostatic interface between chromosome axes and the central region of the SC, facilitated by cation-anion and cation-π interactions. The structural model of phosphorylated SYP-5 was generated using AlphaFold 3. **(D)** Phosphorylation of SYP-5/6 by PLKs reinforces the SC short-arm identity, ensuring accurate segregation of holocentric chromosomes during meiosis I.

Phosphorylation of SYP-5/6 by PLKs begins just prior to meiotic entry (**Fig. 7 A**) and persists along the SC until its disassembly, independent of crossover designation or the PLK-docking site on SYP-1. We envision that SYP-5 and SYP-6 are continuously phosphorylated at the pairing centers (**Fig. 7 B**), where PLK localization and activity are spatially constrained (Roelens et al., 2019; Harper et al., 2011; Labella et al., 2011). As showcased in *hal-2/3* mutants, uncontrolled PLK activity in the nucleoplasm is detrimental and causes precocious loading of SC proteins onto unpaired chromosome axes (Zhang et al., 2012; Roelens et al., 2019). We have now shown that these SC defects in *hal-2* mutants depend on PLK-mediated phosphorylation of the SYP-5/6 C-terminal tails (**Fig. 6 B**), underscoring the importance of spatial control in SYP-5/6 phosphorylation. As homologous chromosomes first come into proximity at the pairing centers, localized phosphorylation of SYP-5/6 by PLKs would permit SC elongation between paired chromosome axes, thereby coupling homolog pairing with the initiation of synapsis (MacQueen et al., 2005).

Our work establishes that the interactions between the SC central region and chromosome axes in *C. elegans* are primarily electrostatic, requiring both PLK-mediated phosphorylation and the adjacent tryptophan residue within the C-terminal tails of SYP-5/6 (**Fig. 7 C**). Disrupting either the phosphorylation event or the tryptophan residues causes severe synapsis defects (**Fig. 3**, **S3, and 5**) and leads to the complete loss of meiotic HORMA domain proteins from polycomplexes in the *rec-8; coh-3 coh-4* mutant (**Fig. 6 A**). We speculate that the tryptophan residues in SYP-5/6 interact with positively charged surfaces on chromosome axes through their polarized π-electron clouds, which can provide substantial binding energy for adhesion (Gallivan and Dougherty, 2000; Gebbie et al., 2017; Waite and Tanzer, 1981). Phosphorylation of the acidic SYP-5/6 C-termini is expected to increase the negative electrostatic potential, thereby enhancing cation-anion interactions with chromosome axes. Furthermore, phosphorylation might induce a conformational change in the disordered C-termini of SYP-5/6, exposing and orienting the π system toward positively charged surfaces on chromosome axes for effective cation-π interactions.

Given the dramatic reduction in affinity for the chromosome axis, it is surprising that some SC stretches still form by late pachytene in our strains harboring mutations in the C-termini of SYP-5/6. This discrepancy may be explained by the larger surface area of meiotic chromosome axes compared to SC aggregates, which can support interactions with the SC central region, albeit at a reduced capacity. Interestingly, the importance of SYP-5/6 phosphorylation becomes most evident after crossover designation, when the SC proteins dissociate from the long arm while becoming enriched on the short arm. Our data indicate that without SYP-5/6 phosphorylation, SC central region proteins prematurely dissociate from chromosome axes, remaining chromosome-bound only through their affinity to crossover-designated sites (**Fig. 3 B**). We speculate that the increased sensitivity to the loss of SYP-5/6 phosphorylation at this stage is due to the remodeling of chromosome axes, which is also dependent on PLK activity (Harper et al., 2011; Sato-Carlton et al., 2018, 2020). Following crossover designation, HTP-1 and HTP-2 are depleted from the short arm in late pachytene (Martinez-Perez et al., 2008), making SYP-5/6 phosphorylation crucial to retain the SC central region. Thus, we hypothesize that prior to their depletion, HTP-1 and HTP-2 on chromosome axes can allow synapsis even in the absence of SYP-5/6 phosphorylation. Consistent with this, loading of SC proteins strictly requires SYP-5/6 phosphorylation throughout the germline in *hal-2/3* mutants, where excessive PLK activity in the nucleoplasm causes premature dissociation of HTP-1/2 from chromosome axes (Roelens et al., 2019; Zhang et al., 2012).

Among the known axis components in *C. elegans*, HIM-3 is positioned closest to the SC central region (Köhler et al., 2017) and promotes synapsis in a dose-dependent manner (Kim et al., 2015; Couteau et al., 2004). Moreover, recent evidence indicates that the recruitment of meiotic HORMA domain proteins to SC polycomplexes in the *rec-8; coh-3 coh-4* mutant depends on the basic residues on HIM-3 (Gordon et al., 2024). Thus, HIM-3 is a likely candidate to interact with the phosphorylated C-terminal tails of SYP-5/6. However, we cannot rule out the possibility that there might be an unidentified protein on chromosome axes that engages in electrostatic interactions with the C-terminal tails of SYP-5/6. Future work is needed to identify the axial counterpart facing the SC central region and to determine how the axis-SC central region interface is dynamically modulated during synapsis.

An unexpected consequence of abolishing SYP-5/6 phosphorylation is the displacement of PLK-2 from chromosome arms during diplotene, which disrupts the asymmetric chromosome remodeling necessary for homolog separation during meiosis I (**Fig. 4**). Our data show that by maintaining the association between the SC central region and chromosome axes, SYP-5/6 phosphorylation helps establish the SC short arm after crossover designation and ensures the proper localization of PLK-2 (**Fig. 7 D**). This mechanism creates a positive feedback loop in which substrate phosphorylation reinforces kinase recruitment, further enhancing the signaling output. We propose that this positive feedback amplifies the signal generated from the crossover-designated site to recruit PLK-2 to the SC (Brandt et al., 2020), thereby driving the robust establishment of the SC short arm and chromosome remodeling, consistent with the recently proposed model (Rodriguez-Reza et al., 2024).

Our findings in *C. elegans* parallel those in budding yeast, where homologous pairing and synapsis depend on crossover recombination (Agarwal and Roeder, 2000; Chua and Roeder, 1998). In budding yeast, synapsis is mediated by the interaction between the transverse filament protein Zip1 and the axis component Red1, which requires small ubiquitin-like modifier (SUMO) and the SUMO-interacting motifs present on these proteins (Hooker and Roeder, 2006; Cheng et al., 2006; Eichinger and Jentsch, 2010). SUMO ligase activity is provided by Zip3, an essential pro-crossover protein that recruits the SUMO-conjugating enzyme Ubc9, thereby initiating homolog synapsis at crossover-designated recombination sites (Hooker and Roeder, 2006; Cheng et al., 2006). Thus, spatially localized signaling at the site of initial homolog contacts, which enables the interaction between the SC central region and chromosome axes, is likely a conserved mechanism for homologous synapsis across diverse organisms.

## MATERIALS AND METHODS

### *C. elegans* strains and CRISPR-mediated genome editing

All strains were maintained on Nematode growth medium (NGM) plates seeded with OP50-1 bacteria at 20°C following standard protocols (Brenner, 1974). N2 Bristol was used as the wild-type strain. **Tables S1 and S3** summarize all mutations and strains used in this study.

To generate point mutations of *syp-5* and *syp-6*, YKM172 (*3xFLAG::syp-5 HA::syp-6 I; meIs8[pie-1p::GFP::cosa-1, unc-119(+)] II*) worms were injected with 16 µM of Cas9 complexed with 16 µM tracrRNA/crRNA oligos (IDT), 5 ng/µl of pCFJ104, 2.5 ng/µl of pCFJ90, and a ssDNA oligo (100 ng/µl) (IDT) as a repair template (**Table S2**). F1 progeny were lysed and genotyped by PCR to detect successful edits. The correct insertion was validated by Sanger sequencing.

### Phospho-antibody production

A synthetic phosphopeptide of SYP-5 flanking S541 (CEEGDADQpSIWGSDD; Biomatik) was coupled to keyhole limpet hemocyanin (KLH) using the Imject™ Maleimide-Activated mcKLH Spin Kit (Thermo Scientific, 77666) and injected into rabbits (Pocono Rabbit Farm & Laboratory) as described in Hurlock et al., 2020. Polyclonal SYP-5 pS541 antibodies were purified by passing the immune serum through SulfoLink Coupling Resins (Thermo Scientific, 44995) coupled to the non-phosphopeptide (CEEGDADQSIWGSDD) and subsequently binding the unbound fraction to phosphopeptide-coupled resins. The specificity of the antibodies was tested through dot blots using both phospho-and non-phosphopeptides, in vitro kinase assays using recombinant 6His-MBP-SYP-5 and GST-PLK-2, and by staining dissected gonads from *syp-5^S541A^ syp-6^T685A^* mutants.

### Protein expression and purification

The full-length open reading frame of SYP-5 was synthesized as a gBlock (IDT) and cloned into pMAL (New England Biolabs) to express Maltose-binding protein (MBP)-tagged SYP-5 with a 6xHis tag at the N-terminus. Protein expression was induced in Rosetta2(DE3)pLysS cells (Novagen) at 15°C with 50 µM IPTG for 16 h. Bacterial cell pellets were resuspended in lysis buffer (PBS, 300 mM NaCl, 50 mM imidazole, 0.5 mM EGTA, 2 mM MgCl2, and 1 mM DTT) containing the cOmplete Protease Inhibitor Cocktail (Sigma 11697498001) and lysed by two freeze/thaw cycles and sonication after lysozyme treatment (0.25 mg/ml) on ice for 30 min. After centrifugation at 15,000 rpm (Beckman, JA-17) for 30 min, the supernatant was loaded onto a HisTrap HP column (Cytiva) and washed with lysis buffer. 6His-MBP-SYP-5 was eluted in elution buffer (PBS, 300 mM NaCl, 500 mM imidazole, 0.5 mM EGTA, 2 mM MgCl2, and 1 mM DTT). 6His-MBP-SYP-5 was further purified by cation exchange using a HiTrap SP HP (Cytiva) column and eluted in a 100 mM to 1M NaCl gradient in 25 mM HEPES, pH 7.5, 2 mM MgCl2, 0.5 mM EGTA, 1 mM DTT. The peak fraction for 6His-MBP-SYP-5 was supplemented with glycerol (20% final) and snap-frozen in liquid nitrogen.

The full-length open reading frame of PLK-2 was amplified from a *C. elegans* cDNA library and cloned into pFastBac1 (Thermo Fisher) with a GST tag at the N-terminus. GST-PLK-2 was expressed for 48 h in *Sf9* cells using the Bac-to-Bac system (Thermo Fisher) and purified over Glutathione Sepharose (Cytiva) using standard protocols. The lysis and purification were conducted in 25 mM HEPES, pH 7.5, 100 mM KCl, 5 mM MgCl_2_, 0.5 mM EGTA, 1 mM DTT, 10 μM MgATP, 0.1% NP-40, and cOmplete Protease Inhibitor Cocktail (Sigma 11697498001).

### In vitro kinase assays and mapping phosphorylation sites

In vitro kinase assays were performed at room temperature in 20 mM HEPES, pH 7.4, 25 mM KCl, 1 mM MgCl_2_, and 1 mM DTT in the presence of 0.2 mM MgATP. 2 µM 6His-MBP-SYP-5 were incubated with 0.1 µM of GST-PLK-2, and the reactions were stopped after 1 hour by boiling in sample buffer and analyzed by Western blot. To map SYP-5 residues phosphorylated by PLK-2 in vitro, phosphorylated 6His-MBP-SYP-5 was cleared using SP3 beads (PreOmics) and digested with trypsin. The resulting peptides were analyzed by MudPIT in the Mass Spectrometry and Proteomics Facility at Johns Hopkins School of Medicine.

### Egg count

L4 hermaphrodites were picked onto individual NGM plates and transferred to new plates every 12 h for 4–5 days. Eggs and hatched L1 larvae were counted immediately after each transfer, and the surviving progeny and males on each plate were counted when the F_1_ generation reached adulthood to quantify the percent egg viability and male self-progeny.

### RNA interference

RNAi was performed by feeding *plk-2(tm1395)* animals with the *Escherichia coli* strain HT115 (DE3), either carrying an Ahringer RNAi library clone targeting *plk-1* or the L4440 empty vector. Bacterial cultures were plated onto RNAi plates (NGM, 25 µg/ml carbenicillin, and 1 mM IPTG), left for 1 h at room temperature to dry, and incubated overnight at 37°C to induce RNA expression. L4 hermaphrodites were picked onto RNAi plates and left for 48 h before dissection for immunofluorescence.

### Immunofluorescence

Hermaphrodite germlines were dissected from 24 h post-L4 adults in egg buffer (25 mM HEPES, pH 7.4, 118 mM NaCl, 48 mM KCl, 2 mM EDTA, 5 mM EGTA, 0.1% Tween-20, and 15 mM NaN_3_), fixed in 2% formaldehyde, frozen in liquid nitrogen, freeze-cracked, and fixed again in ice-cold methanol for 1 minute. For the SYP-5 pS541 staining, fixation was done with 1% paraformaldehyde for 5 minutes, followed by methanol fixation. Slides were rehydrated in PBS with 0.1% Tween-20 and blocked with 1x Roche blocking buffer (Sigma 11096176001) in PBST (0.1% Tween 20) for 45 minutes at room temperature or overnight at 4°C. Incubation with primary antibodies was performed overnight at 4°C at the following concentrations: FLAG (mouse, 1:500; Sigma F1804), SYP-1 (goat, 1:500; MacQueen et al., 2005), SYP-5 (rabbit, 1:1,000; Hurlock et al., 2020), SYP-5 pS541 (rabbit, 1:500; this study), HTP-3 (guinea pig, 1:500; Hurlock et al., 2020), HIM-3 (chicken, 1:500; Hurlock et al., 2020), GFP Booster (1:250; Chromotek gb2AF488), RFP Booster (1:250; Chromotek rba594), Histone H3 pT3 (rabbit, 1:10,000; Cell Signaling Technology 13576S), HTP-1/2 (rabbit, 1:500; Martinez-Perez et al., 2008), HIM-8 (rat, 1:500; Phillips et al., 2005), and phospho-HIM-8/ZIMs (rabbit, 1:1,000; (Kim et al., 2015)). The following secondary antibodies were purchased from Invitrogen or Jackson ImmunoResearch and used at 1:250 dilution: donkey anti-rabbit Alexa Fluor 488, donkey anti-rabbit Alexa Fluor 555, donkey anti-rabbit Alexa Fluor 647, donkey anti-mouse Alexa Fluor 488, donkey anti-mouse Alexa Fluor 555, donkey anti-mouse Alexa Fluor 647, donkey anti-chicken Alexa Fluor 633, donkey anti-guinea pig Alexa Fluor 647, donkey anti-goat Alexa Fluor 647.

Slides were imaged at room temperature with a DeltaVision Elite system (Cytiva) equipped with a 100x oil immersion 1.4 N.A. objective, and a scientific complementary metal-oxide semiconductor camera (Figures 1B-D, 2C, 4A-C, S1B, S2F, S4C, and S5B) or a Leica Thunder Imaging System equipped with a 100x oil immersion 1.4 N.A. objective and a Leica K8 camera (Figures 3, 5C, 6, S2C-D, S3C-D, S4A-B, S5A). For both imaging systems, 3D image stacks were collected at 0.2 μm intervals. Images acquired on the DeltaVision were processed by iterative deconvolution (enhanced ratio, 20 cycles) and projected by the Volume Viewer tool using the SoftWoRx suite (Cytiva). Composite images were assembled and colored in Adobe Photoshop. Images acquired on the Leica Thunder were processed by Large Volume Computational Clearing using the Adaptive Strategy (the feature scale: 341 nm; 92% strength; 30 iterations). Image stacks were projected using the LAS X software (Leica), and the projected images were assembled and colored using Adobe Photoshop.

### Statistical analysis

Statistical analysis was performed using GraphPad Prism 10 software. For comparison of two groups, statistical significance was demonstrated by unpaired *t*-test. For non-bar graphs, error bars show mean and standard deviation. For all graphs, ns indicates no statistical significance, *, *p*<0.05; **, *p*<0.01; ***, *p*<0.001; ****, *p* <0.0001. Sample size is indicated on graphs when greater than one.

### Online supplemental material

Fig. S1 shows validation of the SYP-5 pS541 antibody by dot blot analyses using phospho-and non-phosphopeptides and immunofluorescence in *syp-5^S541A^* and *syp-6^T685A^* mutants. Fig. S2 shows meiotic defects in *syp-5^S541A^*and *syp-6^T685A^* mutants. Fig. S3 shows meiotic defects in *syp-5^S541D^* and *syp-6^T685D^* mutants. Fig. S4 shows additional characterization of SC disassembly in *syp-5^S541D^* and *syp-6^T685D^* and *syp-5^S541A^* and *syp-6^T685A^* mutants. Fig. S5 shows characterization of SC disassembly phenotypes and SYP-5 phosphorylation in *syp-5^W543A^*and *syp-6^W687A^* mutants.

## ACKNOWLEDGEMENTS

We thank Abby Dernburg at the University of California, Berkeley for providing the antibodies, and Bob Cole in the Mass Spectrometry and Proteomics Facility at Johns Hopkins School of Medicine, for mass spectrometry analysis. Some strains were provided by the *Caenorhabditis* Genetics Center (CGC), which is funded by the NIH Office of Research Infrastructure Program (P40OD010440). ALG and MEH were supported by the NIH Predoctoral Fellowships (F31GM150277 and F31HD100090). This work was supported by funding from the National Institutes of Health to YK (R35GM124895).

The authors declare no competing financial interests.

## AUTHOR CONTRIBUTIONS

ALG, MEH, and YK conceived and designed the study. ALG and MEH generated the worm strains. ALG performed most experiments with the assistance of AMG and LYZI. ALG and YK wrote and revised the manuscript.

---

**Figure S1.**
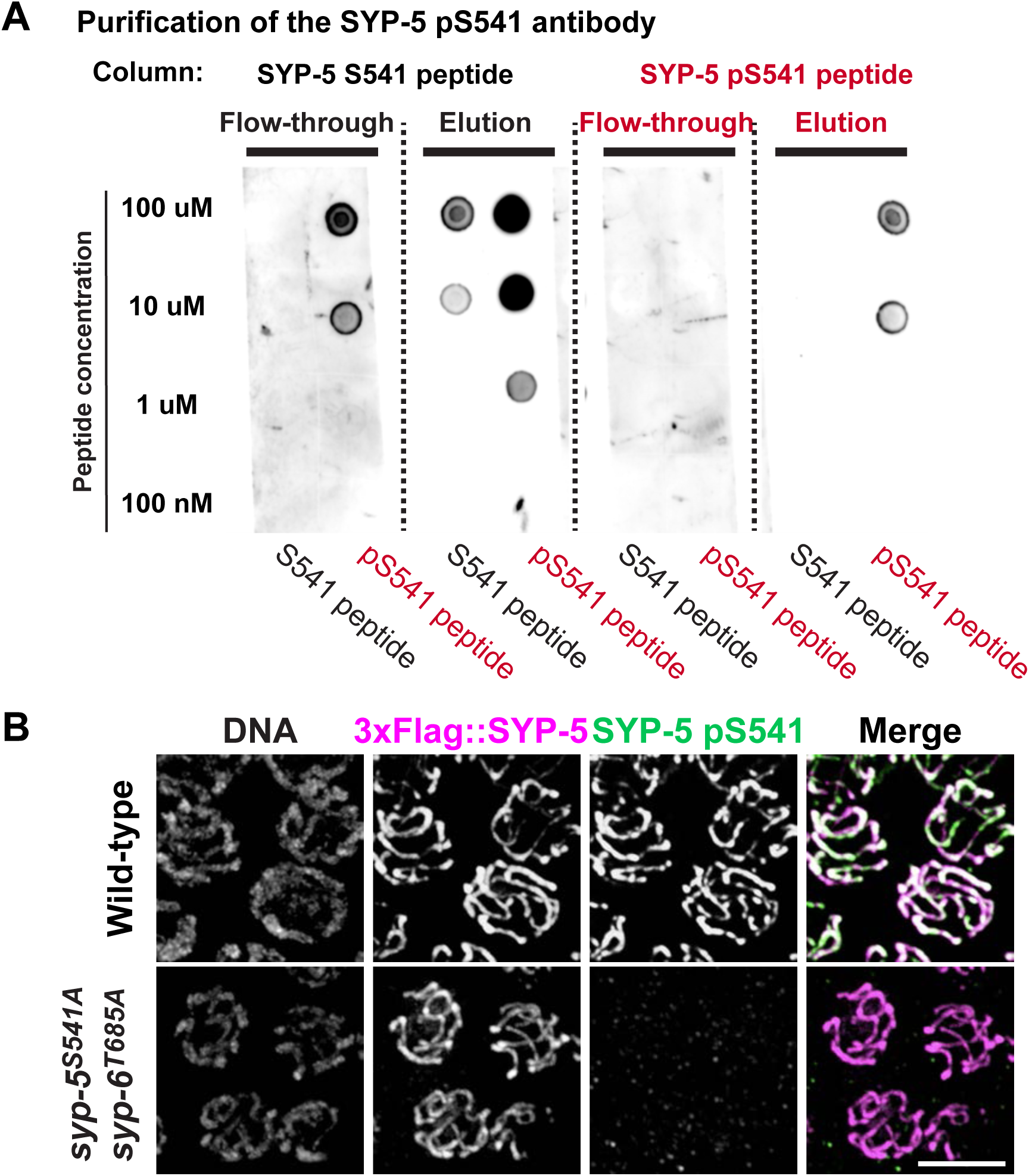
Purification and characterization of the SYP-5 pS541 antibody. **(A)** Immune serum against the SYP-5 phosphopeptide (pS541) was passed through affinity columns coupled to non-phospho (SYP-5 S541) and phosphopeptides (SYP-5 pS541). The unbound (flow-through) and bound (elution) fractions were used in dot blots of serially diluted peptides. **(B)** Immunofluorescence images of pachytene nuclei from wild-type and *syp-5^S541A^ syp-6^T685A^* mutants showing DNA, SYP-5 (magenta), and SYP-5 pS541 (green). Scale bar, 5 µm.

**Figure S2.**
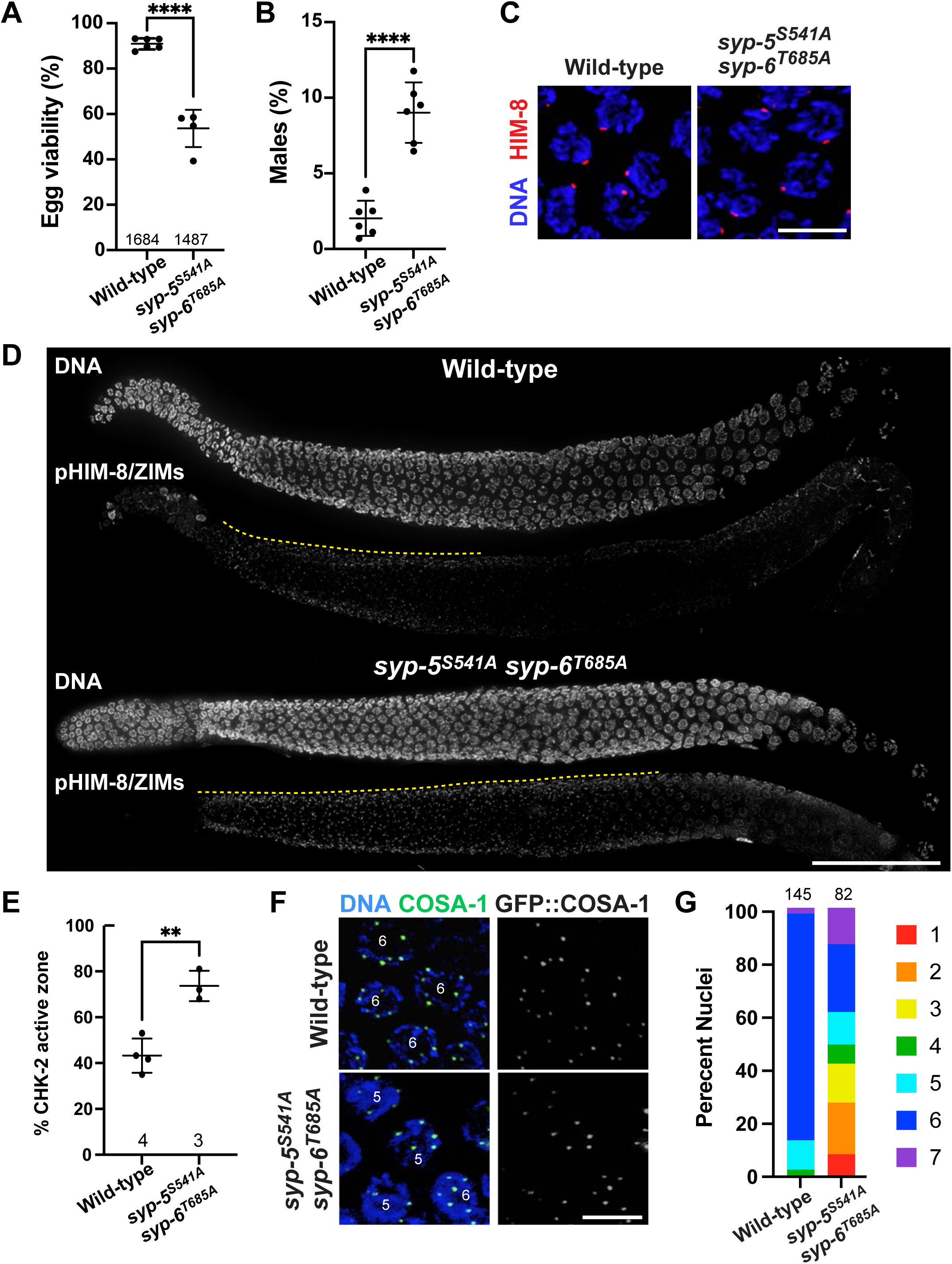
Prolonged CHK-2 activity and abnormal crossover designation in *syp-5^S541A^ syp-6^T685A^* mutants. **(A and B)** Graphs showing the percent viable and male self-progeny from wild-type (n=1684) and *syp-5^S541A^ syp-6^T685A^* mutant (n=1487) hermaphrodites. ****, p<0.0001 by unpaired t-test. **(C)** Immunofluorescence images from wild-type and *syp-5^S541A^ syp-6^T685A^* mutants stained for DNA (blue) and HIM-8 (red). Scale bar, 5 µm. **(D)** Dissected gonads from wild-type and *syp-5^S541A^ syp-6^T685A^* mutants were stained for DNA and phospho-HIM-8/ZIMs (CHK-2 substrate). Composite projection immunofluorescence images are shown. The CHK-2 active zone is marked by yellow dotted lines. Scale bar, 50 µm. **(E)** Graph showing the CHK-2 active zone in wild-type and *syp-5^S541A^ syp-6^T685A^* mutants, shown as a percent gonad length of the pHIM-8/ZIM-positive region relative to the region from meiotic entry to pachytene exit. The numbers of gonads scored are also indicated. **, p<0.05 by unpaired t-test. **(F)** Immunofluorescence images of pachytene nuclei from wild-type and *syp-5^S541A^ syp-6^T685A^* animals stained for DNA (blue) and GFP::COSA-1 (green). Scale bar, 5 µm. **(G)** Graph showing the quantification of GFP::COSA-1 foci per nucleus. The numbers of nuclei scored are indicated on the top.

**Figure S3.**
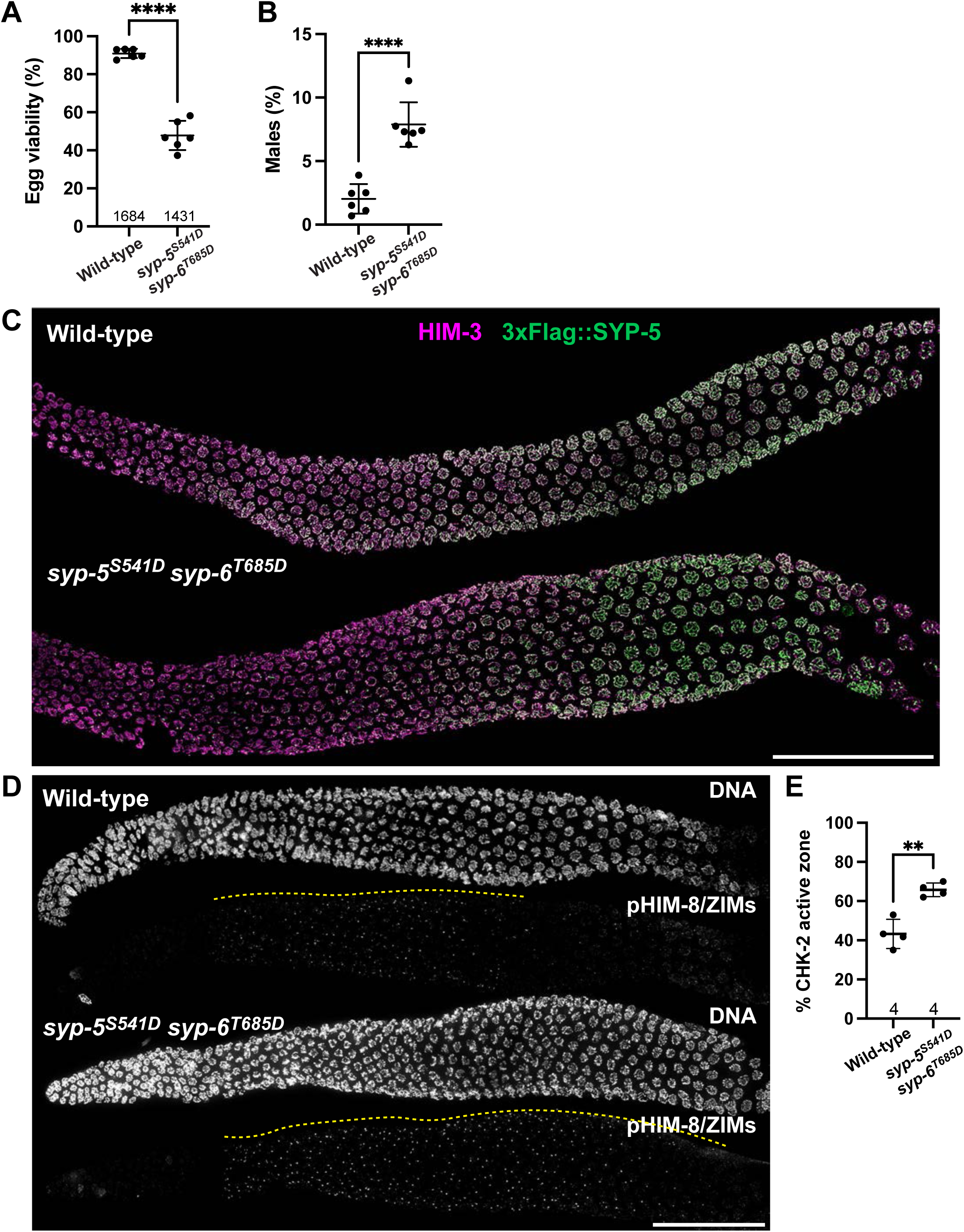
Phospho-mimetic mutations at SYP-5 S541 and SYP-6 T685 cause defects in SC assembly. **(A-B)** Graph showing the percent viable and male self-progeny from the wild type (n=1684) and *syp-5^S541D^ syp-6^T685D^* mutants (n=1431) hermaphrodites. ****, p<0.0001 by unpaired t-test. **(C)** Full-length gonads dissected from wild-type and *syp-5^S541D^ syp-6^T685D^* animals were stained for HIM-3 (magenta) and 3xFlag::SYP-5 (green). Composite projection images are shown. Scale bar, 50 µm. **(D)** Composite immunofluorescence images of dissected gonads from wild-type and *syp-5^S541D^ syp-6^T685D^* animals with DNA and phospho-HIM-8/ZIMs (CHK-2 substrate) staining are shown. Yellow dotted lines mark the CHK-2 active zone. Scale bar, 50 µm. **(E)** Graph showing the CHK-2 active zone in the wild type and *syp-5^S541A^ syp-6^T685A^*mutants, shown as a percent gonad length of the pHIM-8/ZIM-positive region relative to the region from meiotic entry to pachytene exit. The numbers of gonads scored are also indicated. **, p=0.0015 by unpaired t-test.

**Figure S4.**
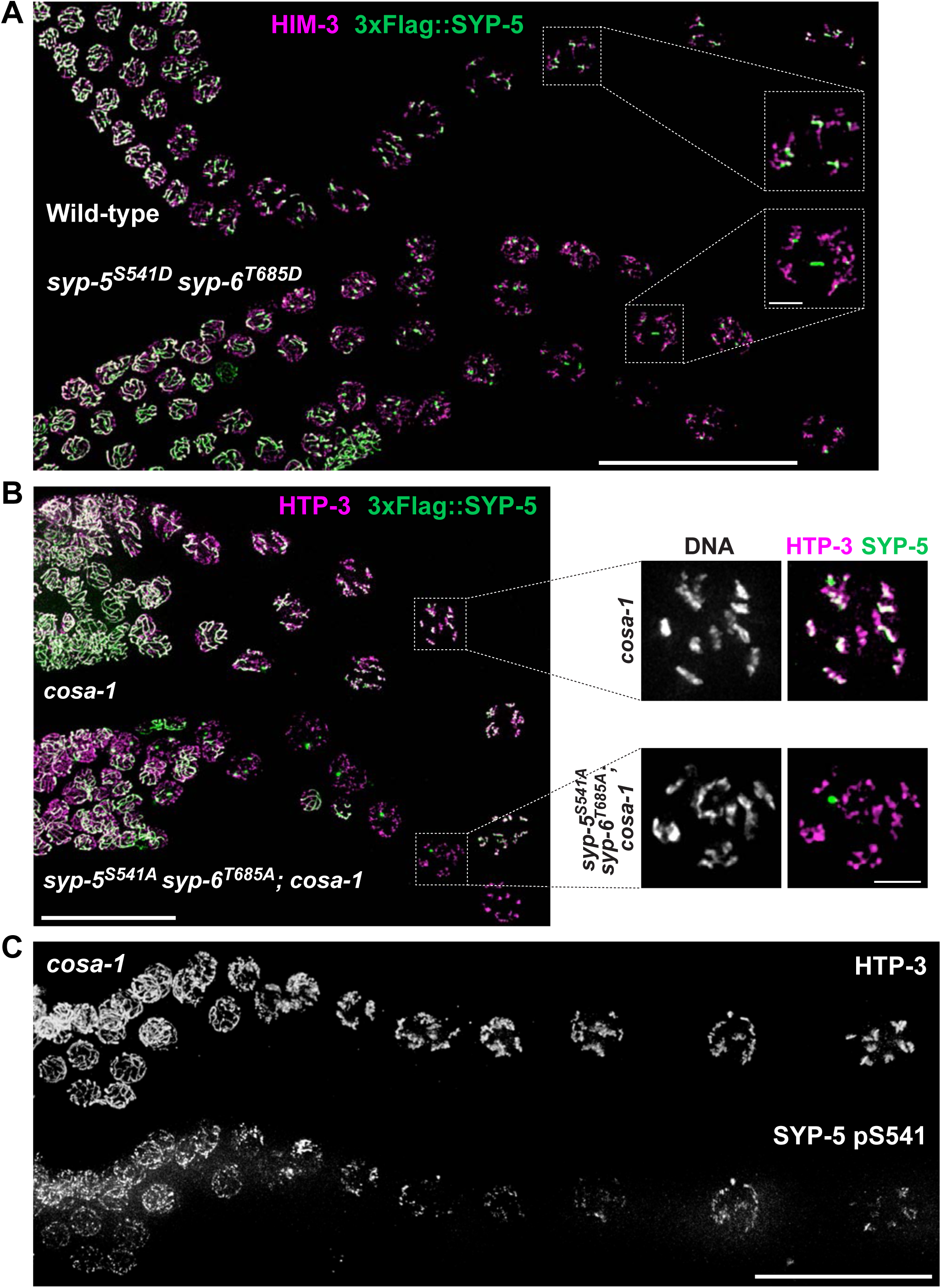
Additional characterization of SYP-5/6 phosphorylation and SC disassembly defects in *syp-5/6* phosphomutants. **(A)** Immunofluorescence images of diplotene nuclei from wild-type and *syp-5^S541D^ syp-6^T685D^* mutant animals stained for HIM-3 (magenta) and SYP-5 (green). Scale bar, 25 µm. The inset shows the zoomed-in view of the boxed regions. Scale bar, 1 µm. **(B)** Immunofluorescence images of diplotene nuclei from *cosa-1* and *syp-5^S541D^ syp-6^T685D^; cosa-1* mutants stained for HTP-3 (magenta) and SYP-5 (green). Scale bar, 25 µm. The inset shows the zoomed-in view of the boxed regions. Scale bar, 1 µm. **(C)** Immunofluorescence images of diplotene nuclei from *cosa-1(tm3298)* mutants. HTP-3 and SYP-5 pS541 staining are shown. Scale bar, 25 µm.

**Figure S5.**
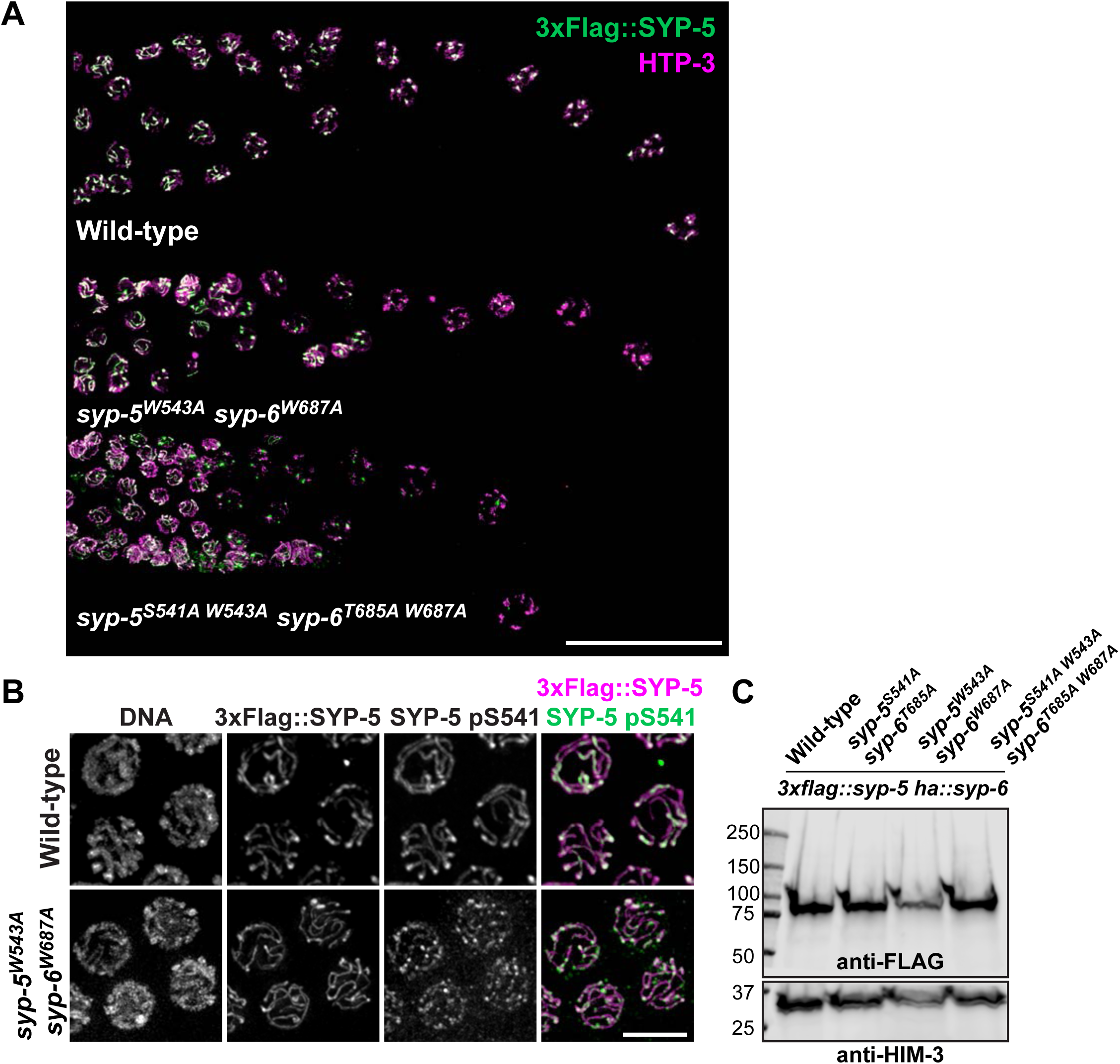
Characterization of SC disassembly and SYP-5 phosphorylation in *syp-5^W543A^ syp-6^W687A^* mutants. **(A)** Immunofluorescence images of diplotene nuclei from wild-type and *syp-5^W543A^ syp-6^W687A^* mutant animals stained for 3xFlag::SYP-5 (green) and HTP-3 (magenta). Scale bar, 25 µm. **(B)** Immunofluorescence images of pachytene nuclei from wild-type and *syp-5^W543A^ syp-6^W687A^* animals showing DNA, 3Flag::SYP-5 (magenta), and SYP-5 pS541 (green) staining. Scale bar, 5 µm. **(C)** Western blot of worm lysates from the indicated genotypes probed against the Flag tag. HIM-3 was used as a loading control.

**TABLE S1.** Alleles generated in this study.

| Allele | Genotype | Information about mutagenesis |
| --- | --- | --- |
| *kim18* | *syp-5(kim18[3xFLAG::syp-5(S541A)]) I* | Generated using CRISPR in YKM172 (3xFLAG::syp-5 HA::syp-6) |
| *kim19* | *syp-6(kim19[HA::syp-6(T685A)]) I* | Generated using CRISPR in YKM172 (3xFLAG::syp-5 HA::syp-6) |
| *kim79* | *syp-5(kim79[3xFLAG::syp-5(S541D)]) I* | Generated using CRISPR in YKM172 (3xFLAG::syp-5 HA::syp-6) |
| *kim80* | *syp-6(kim19[HA::syp-6(T685D)]) I* | Generated using CRISPR in YKM172 (3xFLAG::syp-5 HA::syp-6) |
| *kim82* | *syp-5(kim82[3xFLAG::syp-5(W543A)]) I* | Generated using CRISPR in YKM172 (3xFLAG::syp-5 HA::syp-6) |
| *kim83* | *syp-6(kim83[HA::syp-6(W687A)]) I* | Generated using CRISPR in YKM172 (3xFLAG::syp-5 HA::syp-6) |
| *kim84* | *syp-5(kim84[3xFLAG::syp-5(S541D W543A)]) I* | Generated using CRISPR in YKM353 (3xFlag::syp-5<sup>S541A</sup> HA::syp-6<sup>T685A</sup>) |
| *kim85* | *syp-6(kim85[HA::syp-6(T685A W687A)]) I* | Generated using CRISPR in YKM353 (3xFlag::syp-5<sup>S541A</sup> HA::syp-6<sup>T685A</sup>) |

**TABLE S2.**
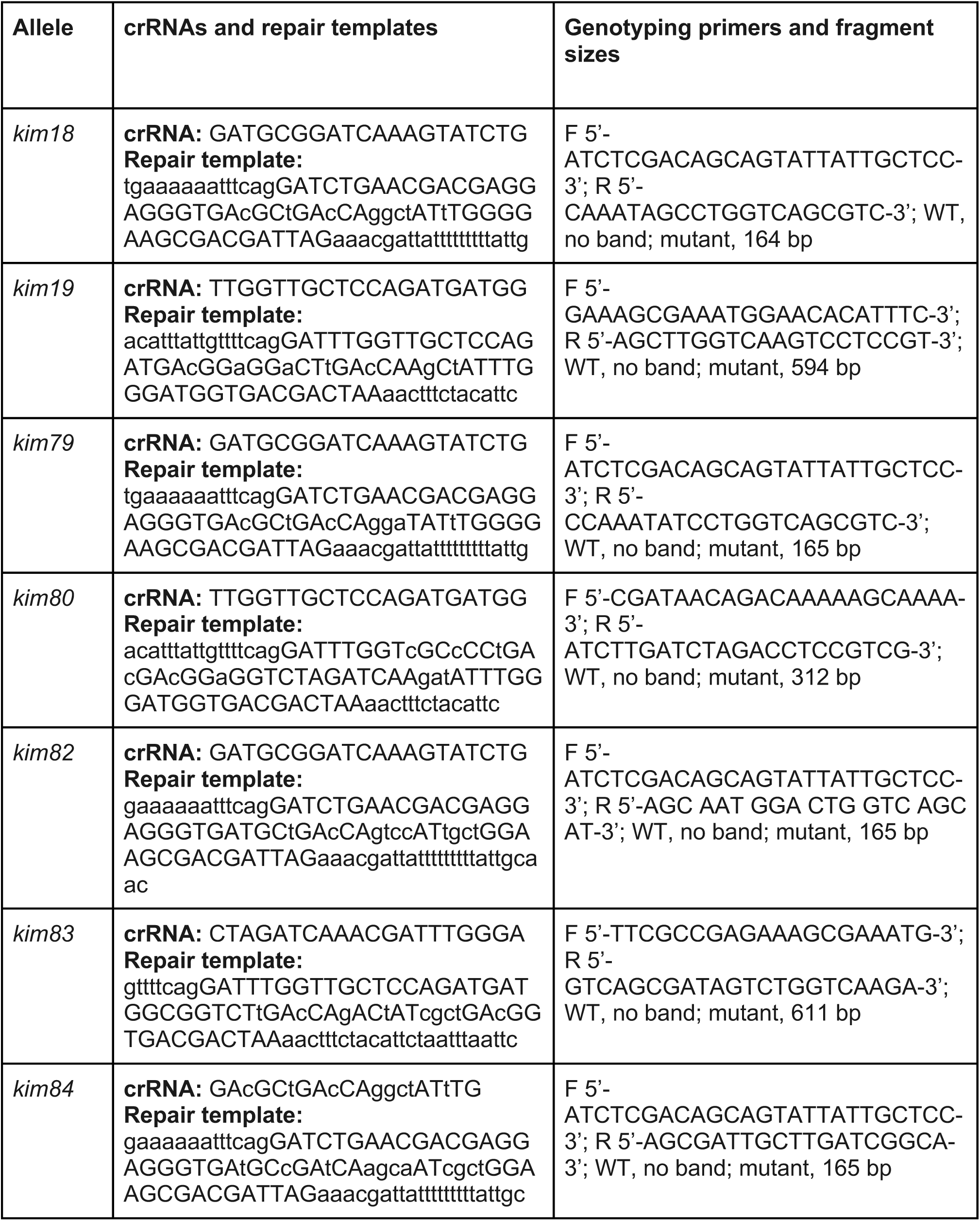
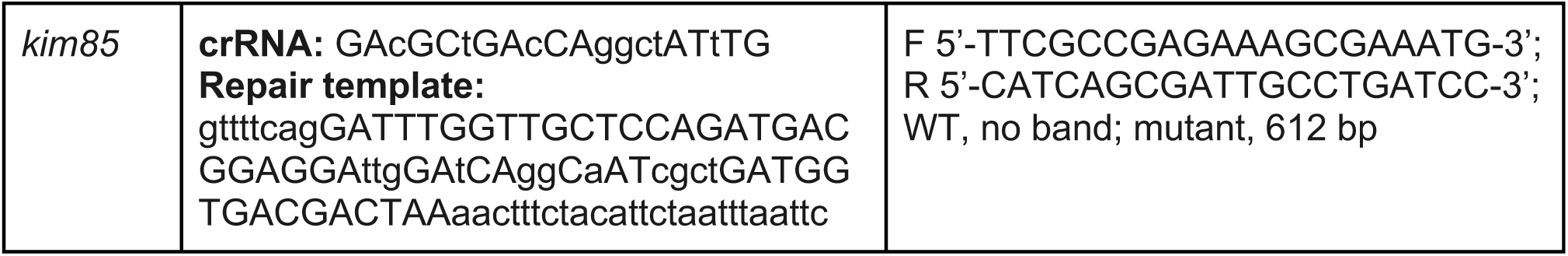
crRNAs, repair templates, and genotyping primers for mutant alleles generated in this study.

**TABLE S3.**
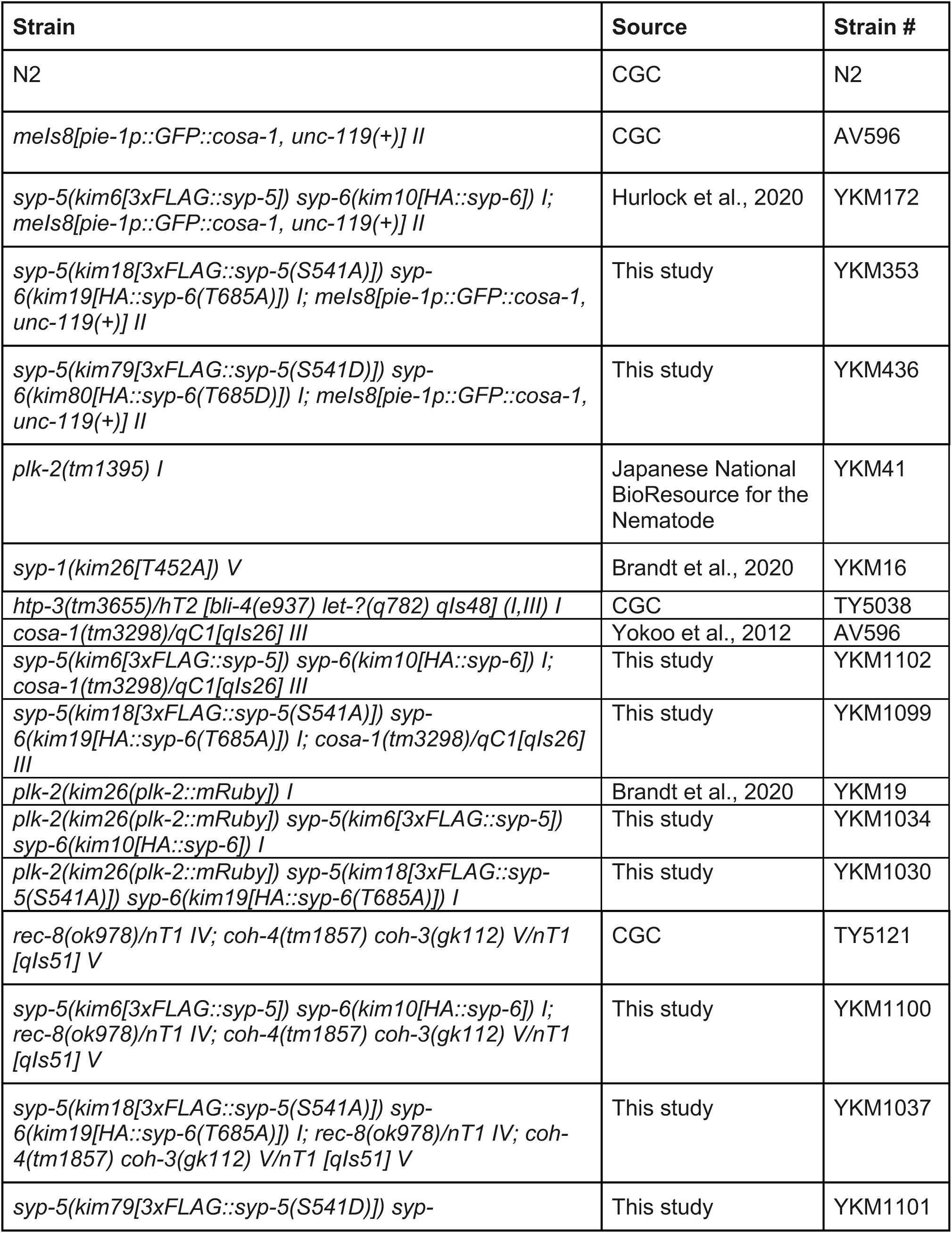
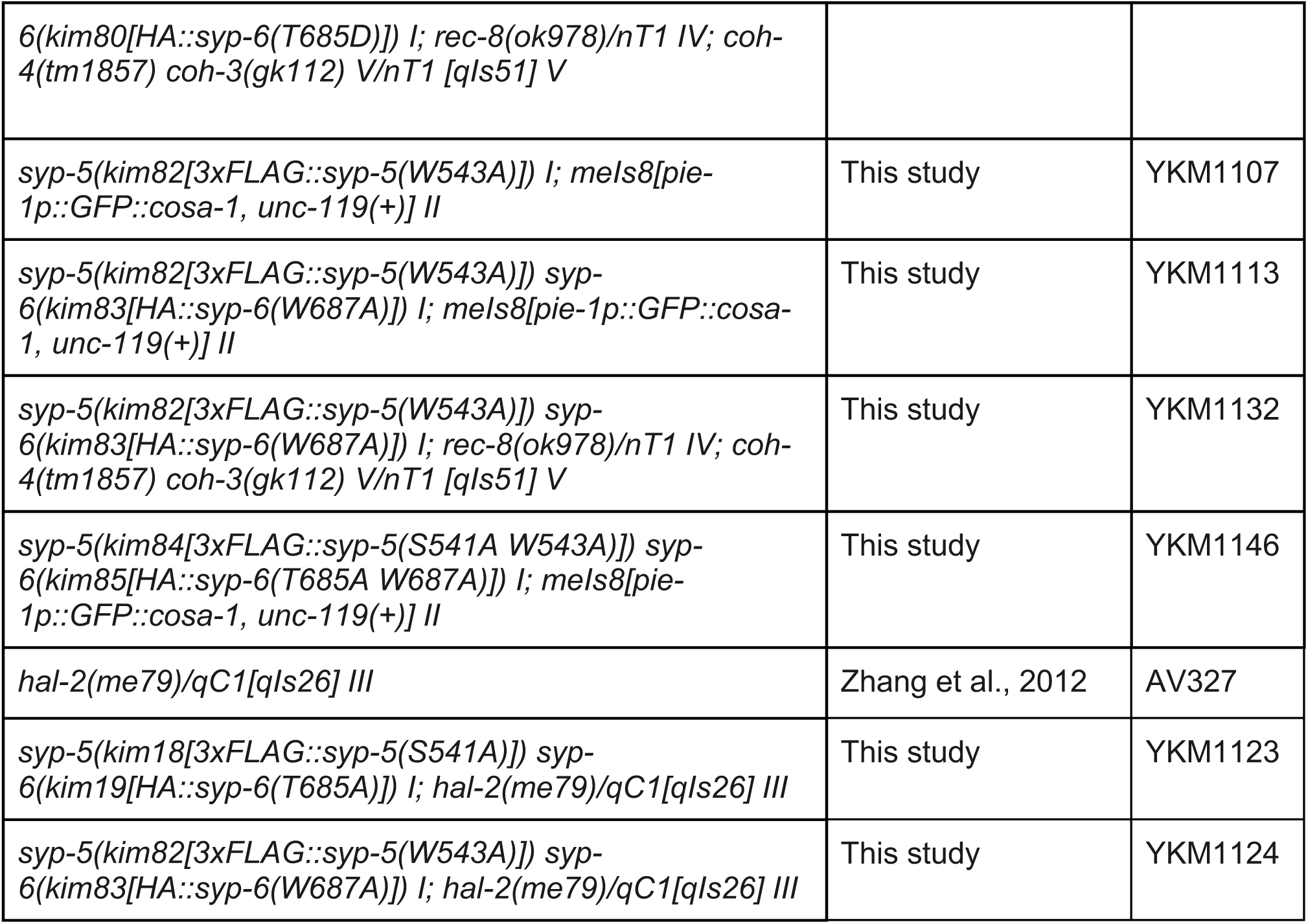
Strains used in this study.

